# AtpΘ is an inhibitor of F_0_F_1_ ATP synthase to arrest ATP hydrolysis during low-energy conditions in cyanobacteria

**DOI:** 10.1101/2021.07.30.454416

**Authors:** Kuo Song, Desirée Baumgartner, Martin Hagemann, Alicia M. Muro-Pastor, Sandra Maaß, Dörte Becher, Wolfgang R. Hess

## Abstract

Biological processes in all living cells are powered by ATP, a nearly universal molecule of energy transfer. ATP synthases produce ATP utilizing proton gradients that are usually generated by either respiration or photosynthesis. However, cyanobacteria are unique in combining photosynthetic and respiratory electron transport chains in the same membrane system, the thylakoids. How cyanobacteria prevent the futile reverse operation of ATP synthase under unfavorable conditions pumping protons while hydrolyzing ATP is mostly unclear. Here, we provide evidence that the small protein AtpΘ, which is widely conserved in cyanobacteria, is mainly fulfilling this task. The expression of AtpΘ becomes induced under conditions such as darkness or heat shock, which can lead to a weakening of the proton gradient. Translational fusions of AtpΘ to the green fluorescent protein revealed targeting to the thylakoid membrane. Immunoprecipitation assays followed by mass spectrometry and far Western blots identified subunits of ATP synthase as interacting partners of AtpΘ. ATP hydrolysis assays with isolated membrane fractions as well as purified ATP synthase complexes demonstrated that AtpΘ inhibits ATPase activity in a dose-dependent manner similar to the F_0_F_1_-ATP synthase inhibitor N,N-dicyclohexylcarbodimide. The results show that, even in a well-investigated process, crucial new players can be discovered if small proteins are taken into consideration and indicate that ATP synthase activity can be controlled in surprisingly different ways.

## Introduction

ATP synthases of the F_0_F_1_ type are multisubunit protein complexes anchored to membranes that convert proton (or sodium ion) gradients into chemical energy in the form of ATP^1^. Proton gradients are established by divergent processes, such as respiratory electron transport in mitochondria or photosynthetic electron transport in chloroplasts. Mitochondria and chloroplasts originate from the endosymbiotic uptake of an α-proteobacterium and a cyanobacterium, respectively^2–7^. Therefore, it is not surprising that F_0_F_1_-ATP synthases share close functional and structural similarities among eukaryotes and bacteria.

Under conditions weakening the proton gradient, ATP synthases can operate backwards, pumping protons while hydrolyzing ATP. Therefore, different regulatory mechanisms have evolved to stop the futile reverse reaction. Mitochondrial ATP synthases employ small peptides for inhibition, one, designated inhibitory factor 1 (IF1), in mammals^8, 9^ and three, called IF1, STF1 and STF2, in yeast^10, 11^. IF1 inhibits the ATPase activity of mitochondrial ATP synthase under conditions when the membrane potential collapses, e.g., during anoxia in cancer cells^12^. In bacteria, some regulatory factors of ATP synthase are known as well, such as the ζ subunit in *Paracoccus denitrificans* and related α-proteobacteria^13^, but IF1, as a representative of the class of alpha-helical basic peptide inhibitors in eukaryotes, has no homologs among prokaryotes.

Plant chloroplasts, in contrast, use a different mechanism to inhibit the hydrolysis activity of ATP synthase. Here, the γ subunit encoded by *atpC* responds to redox signals, thereby preventing the back reaction of ATP synthase when the photosynthetic proton gradient ceases, particularly during the night^14^. The *atpC* gene and the encoded γ subunit in chloroplasts are very similar to their homologs from cyanobacteria, consistent with the endosymbiotic origin of chloroplast ATP synthase from an ancient cyanobacterium^15^. The chloroplast γ subunit, however, possesses a short insertion of nine extra amino acids (–EICDINGXC–), including two cysteine residues^16^ that can form a disulfide bond under oxidizing conditions, which entirely blocks rotation and prevents ATP hydrolysis^1^. Upon illumination, the chloroplasts become reduced, and the disulfide bridge in the γ subunit opens, which activates ATP synthase because the γ subunit can rotate freely. The respective nine-amino-acid insertion in chloroplast γ subunits is strictly conserved in plants but missing from any of the homologs in cyanobacteria^17^. In contrast to chloroplasts, in cyanobacteria, photosynthetic and respiratory electron transport chains are both located in the same membrane system, the thylakoids, and even share some components^18^. Therefore, cyanobacteria cannot shut down ATP synthase as strictly as plant chloroplasts during the dark phase, since both the photosynthetic and respiratory electron chains generate proton gradients at the thylakoid membranes during day and night, respectively, which are used by the same ATP synthase for the generation of ATP^19^. Hence, the cyanobacterial ATP synthase complexes cannot be controlled by the same redox-sensitive mechanism as operating in the chloroplast.

Nevertheless, several mechanisms have been identified for the regulation of ATP synthase activity in cyanobacteria, the ADP-mediated inhibition that relies on the γ subunit^20^ and ε subunit-mediated inhibition^21^. These findings provided hints that also mechanisms to prevent wasteful ATP hydrolysis activity of ATP synthase might exist in cyanobacteria.

Here, we provide evidence that a small protein previously called Norf1 (for novel ORF1) acts as ATP synthase regulator in cyanobacteria. Norf1 was initially discovered in the model cyanobacterium *Synechocystis* sp. PCC 6803 (*Synechocystis* 6803) based on the detection of its mRNA in transcriptomic datasets^22, 23^. *Synechocystis* 6803 Norf1 comprises 48 amino acids, and its expression was confirmed at the protein level by Western blot analyses^24^. The *norf1* mRNA level was found to increase dramatically after the transfer of cultures into darkness^22^. Darkness-stimulated gene expression is very unusual in cyanobacteria, that base their physiology on light-dependent oxygenic photosynthesis. In *Synechocystis* 6803, only 62 out of a total of 4,091 experimentally defined transcriptional units exhibited maximum expression in the dark^22^. Therefore, it appeared elusive why a free-standing gene encoding a small protein of just 48 amino acids would be regulated in this way and make its transcript the mRNA with the highest absolute read count after 12 h in darkness^22^.

To elucidate Norf1 function, we scrutinized its expression here in more detail, investigated mutant strains and identified interacting proteins. Norf1 is a soluble protein, but membrane fractionation experiments and fusions to GFP showed targeting to the thylakoid membrane. Immunoprecipitation followed by mass spectrometry and far Western blot suggested specific interactions with subunits of the ATP synthase complex. Finally, measurements of ATP hydrolysis in isolated membrane fractions, and purified ATP synthase complexes revealed that Norf1 is recruited during unfavorable conditions as an inhibitory subunit that prevents the hydrolysis of ATP. These findings prompted us to rename Norf1 and its gene to AtpΘ for the cyanobacterial <u>ATP</u> synthase inhibi<u>T</u>ory factor (gene *atpT*).

## Results

### Genes encoding homologs of AtpΘ are widely distributed throughout the cyanobacterial phylum

The 48 amino acid sequence of the previously identified *Synechocystis* 6803 Norf1 protein^24^, here renamed AtpΘ, was used to search for homologs, resulting in the identification of highly similar proteins in 318 available cyanobacterial genomes, including all finished genomes and some of the permanent draft genomes (**Figure 1**). The occurrence, sequence and predicted isoelectric points of AtpΘ homologs are given in **Table S1**. These homologs were predicted based on quite short amino acid sequences; therefore, we cannot rule out that the list includes some false positives or that some homologs might have been missed. While most cyanobacterial genomes (228/318) possess a single *atpT* gene, we also identified 88 genomes with two and two genomes with three putative homologs (**Figure S1A**). Putative *atpT* homologs were not detected outside the cyanobacterial phylum, but homologs were found in two *Gloeobacter* species considered to represent the most ancestral clade^25^, pointing at an early and stable acquisition of *atpT* in the cyanobacterial radiation (**Figure 1**). Most of the genomes containing two homologs are relatively large (median 6.23 Mb) and belong mainly to the genera *Fischerella, Calothrix, Scytonema* and *Nostoc*. The different copies in one strain are not identical, making their origin from recent gene duplications unlikely. The majority of the putative homologs are between 39 and 70 amino acids in length (**Figure S1B**), except those in *Halomicronema hongdechloris* C2206 and *Pseudanabaena* sp. PCC 7367 with 94 and 82 amino acids, respectively. However, the homolog in *Pseudanabaena* sp. PCC 7367 exhibits pronounced sequence similarity only within its central and C-terminal residues, potentially being translated from an internal start codon (marked in **Table S1** in red) and yielding a peptide of 51 residues.

**Figure 1.**
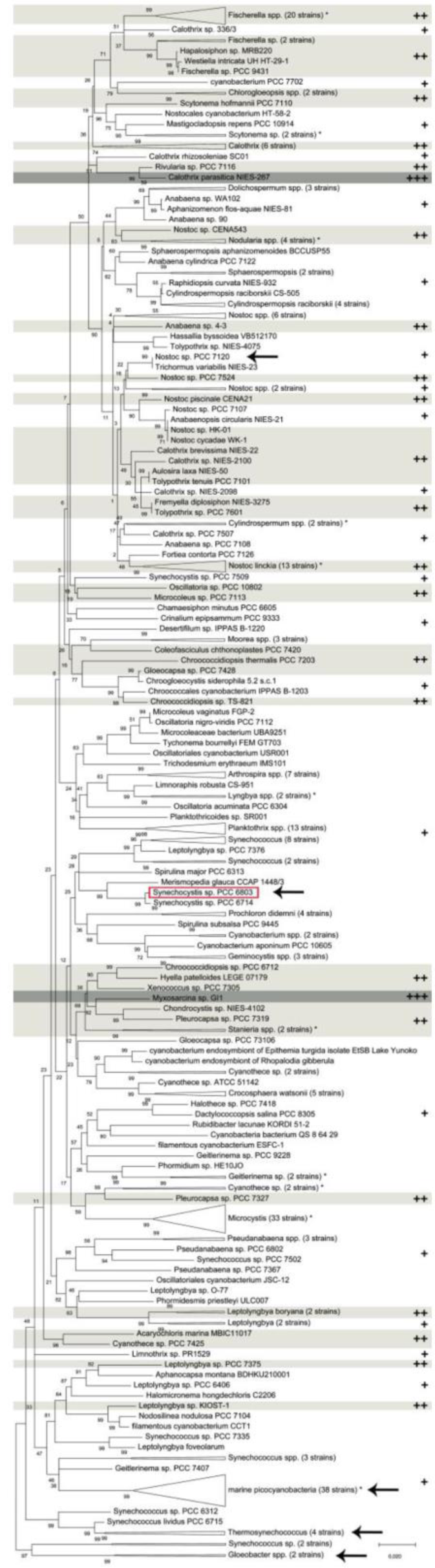
Distribution and numbers of *atpT* genes throughout the cyanobacterial phylum. Phylogenetic tree of cyanobacteria based on 16S rRNA sequences (SILVA database^43^ constructed in MEGAX^44^ using the minimum evolution method^45^). The number of individual strains is given in brackets if several strains were joined at one branch (e.g., 33 strains for *Microcystis*), marine picocyanobacteria consisting of *Prochlorococcus* and marine *Synechococcus*. The numbers of putative *atpT* homologs in each strain are indicated (+, one; ++, two; +++, three homologs) and additionally highlighted in shades of gray if more than one. Single deviations within clusters of strains joined at one branch are labeled by asterisks (e.g., among the 20 *Fischerella* spp. strains in the uppermost cluster is one strain with one homolog, while all others have two). Species selected for experimental analyses in this study are labeled by arrows, and the location of the *Synechocystis* 6803 model strain is additionally highlighted by a red box. The optimal tree with the sum of branch length = 5.06878059 is shown. The percentage of replicate trees in which the associated taxa clustered together in the bootstrap test (500 replicates) is shown next to the branches^46^. The tree is drawn to scale, with branch lengths in the same units as the units of the evolutionary distances used to infer the phylogenetic tree. The analysis involved 318 nucleotide sequences. All ambiguous positions were removed for each sequence pair (pairwise deletion option). There were a total of 1753 positions in the final dataset. The sequences of all potential AtpΘ homologs are given in **Table S1**.

AtpΘ homologs are predicted to be soluble proteins lacking transmembrane helices. Sequence comparison of selected AtpΘ homologs covering strains from all identified larger phylogenetic clusters among cyanobacteria showed quite different sequences, with only 7 widely conserved residues (**Figure S1C**). These residues include aromatic residues at positions 13 and 22, negatively charged residues at positions 16 and 27 and a conserved proline at position 30 with regard to the *Synechocystis* 6803 protein. This divergence is also reflected in the isoelectric points (IPs), which were predicted to range from acidic values (AtpΘ in *Synechocystis* 6803 and *Microcystis*) to very alkaline values (>11) for AtpΘ from thermophilic strains (**Table S1**).

### Energy supply and proton gradient integrity impact *atpT* transcription

Northern blot experiments showed that the *atpT* transcript level increased within 10 min after transfer to darkness, rapidly reaching maximum values 30 min after transfer and declined only marginally at the latest time point (**Figure 2A**). The addition of 10 mM glucose neutralized the strong darkness-induced activation of gene expression (**Figure 2A**), suggesting that the stimulation of *atpT* transcript accumulation in the dark is connected to the energy supply for respiration. Based on these results, we chose an incubation time of 6 h in darkness for subsequent experiments. High *atpT* expression was also previously associated with transfer to darkness or low light conditions, entering stationary phase or heat shock^22, 24^. We reasoned that all these conditions compromise photosynthetic activity and may affect the cellular redox status. Therefore, we tested additional conditions that interfere with the proton gradient or the electron transfer chain. Indeed, the parallel presence of the uncoupler carbonyl cyanide *m*-chlorophenyl hydrazone (CCCP)^26^ or of the electron transport inhibitor 2,5-dibromo-3-methyl-6-isopropyl-*p*-benzoquinone (DBMIB)^27^ restored the high transcript accumulation in the dark despite the addition of glucose (**Figure 2B**). These results indicated that it was not the lack of light *per se* that triggered *atpT* expression. Instead, the enhanced respiration fostered by the addition of glucose led to the suppression of the dark-induced increase in transcript accumulation, while CCCP or DBMIB lifted this suppression. We conclude that it was the potentially low capacity for ATP synthesis due to a diminished or absent proton gradient that triggered high *atpT* expression.

**Figure 2.**
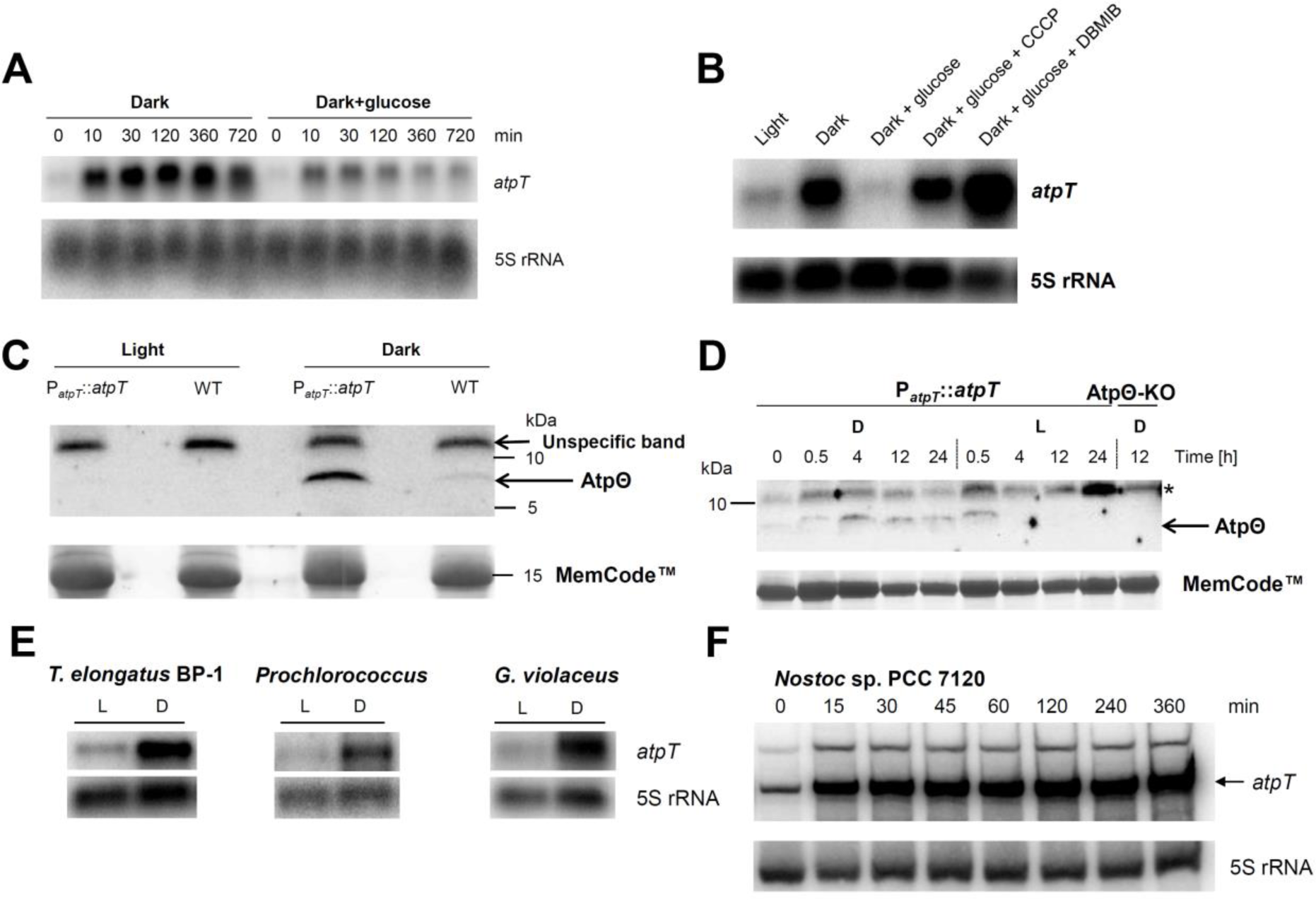
Expression of *atpT* is stimulated by low energy, uncoupling or inhibition of electron transfer. **(A)** Time course of *atpT* mRNA accumulation in the dark in the presence or absence of glucose (10 mM). Exponentially growing *Synechocystis* 6803 wild-type (WT) cells were harvested at the indicated time points before and after transfer to darkness. Northern hybridization was carried out after separation and blotting of 1 μg of total RNA with a ^32^P-labeled, single-stranded transcript probe specifically recognizing *atpT*. **(B)** *atpT* mRNA accumulation after 6 h under the indicated conditions and treatments. CCCP and DBMIB were added to final concentrations of 10 µM and 100 µM, respectively. **(C)** Western blot experiment for the detection of native AtpΘ. Protein levels were compared in *Synechocystis* 6803 WT cells and in cultures carrying the P*atpT*::*atpT* construct in which the untagged *atpT* gene was overexpressed from its native promoter on the plasmid vector pVZ322 in addition to the native gene copy. Identical amounts of 150 µg total protein were separated by Tricine SDS-PAGE^47^ and probed with anti-AtpΘ serum after transfer to nitrocellulose membrane. Precision Plus Protein™ DualXtra (2-250 kDa, Bio-Rad) was used as molecular mass marker. The same membrane was stained with MemCode™ as a loading control. **(D)** AtpΘ expression under changing light conditions. Samples for protein extraction were collected at the indicated time points. Approximately 150 µg (calculated according to Direct Detect™ Spectrometer measurements) of protein samples was separated. PageRuler™ Prestained Protein Ladder (10–170 kDa, Fermentas) was used as molecular mass marker. MemCode™ staining served as a loading control. D, darkness, L, standard light (∼40 µmol photons m^-2^ s^-1^), AtpΘ-KO, AtpΘ knockout (only last lane). In panels **(C)** and **(D)**, the position of untagged AtpΘ is indicated, * indicates a strong cross-reacting band. **(E)** Northern analysis of potential *atpT* homologs in *Thermosynechococcus elongatus* BP-1, *Prochlorococcus* sp. MED4, and *Gloeobacter violaceus* PCC 7421. For each sample, 5 μg of total RNA was loaded. L, strains were cultured in constant light; D, light-cultured strains were incubated in darkness for 6 h. **(F)** Time course of *atpT* mRNA accumulation in *Nostoc* 7120 in cultures transferred from light to darkness for the indicated times. In panels (A), (B), (E) and (F) the respective 5S rRNA was hybridized as a loading control.

To evaluate the accumulation of the AtpΘ protein, a specific antibody was raised that detected a faint band with an apparent molecular mass of 8 kDa in samples from *Synechocystis* 6803 wild-type cultures grown in the dark but not in the light (**Figure 2C**). Expression of AtpΘ under the control of its native promoter from plasmid vector pVZ322 enhanced the detected band more than twofold, caused by the higher copy number of the plasmid-located gene. Western blot analysis also showed that AtpΘ started to accumulate 0.5 h after transfer to darkness and continued to become more abundant over a time period of 4 h, after which it remained at approximately the same level; transfer of the cultures back into light led to the disappearance of the AtpΘ signal within less than 4 h (**Figure 2D**). Thus, the time course of AtpΘ protein accumulation after transfer of cultures into darkness closely followed the time course of mRNA accumulation.

The inducibility by transfer into darkness might be characteristic of *atpT* expression and might support the identification of putative homologs in different species. We chose four species that are phylogenetically distant from *Synechocystis* 6803 (**Figure 1**). *Gloeobacter violaceus* PCC 7421 represents an early-branching species that lacks thylakoid membranes^28^. *Thermosynechococcus elongatus* BP-1 belongs to a clade of unicellular thermophilic strains, while *Prochlorococcus* sp. MED4 is a laboratory isolate representing the vast marine picocyanobacterial genus *Prochlorococcus*^29^. Finally, *Nostoc* sp. PCC 7120 (*Nostoc* 7120) is a model strain for the group of heterocyst-differentiating and N2-fixing multicellular cyanobacteria. The predicted AtpΘ homologs share as little as 12.5% (*Prochlorococcus* sp. MED4), 20.8% (*G. violaceus* PCC 7421), 33.3 and 41.4% (*T. elongatus* BP-1 and *Nostoc* 7120) identical amino acids with the *Synechocystis* 6803 protein. The results of Northern hybridizations showed that the predicted *atpT* homologs in all four strains were expressed at higher levels after 6 h in darkness than under light conditions (**Figures 2E** and **2F**). These findings reinforced the idea that these genes, identified only on the basis of sequence searches, might be orthologs of the *atpT* gene in *Synechocystis* 6803.

### AtpΘ localizes in *Synechocystis* 6803 to soluble and membrane-enriched protein fractions

Strain P_*atpT*_::*atpT*-3xFLAG was used to localize AtpΘ within soluble or membrane-enriched protein fractions. To verify the specificity of the Flag antibody, we also analyzed strains P_*atpT*_::*atpT* (negative control) and P_*petJ*_::3xFLAG-sf*gfp* (positive control). *Synechocystis* 6803 extracts from dark- and light-grown cultures were separated by centrifugation into membrane and soluble fractions and analyzed by Western blotting. FLAG-tagged proteins were detected in the respective lysates, while no signal was obtained for the negative control. AtpΘ was partitioned approximately equally between the soluble and membrane fractions, while FLAG-tagged sfGFP was restricted to the soluble fraction (**Figure 3A**). The FLAG tag stabilized the AtpΘ protein, since the FLAG-tagged version could be detected in samples from cultures kept under continuous light for 12 h, very different from the native form (**Figures 2C** and **2D**). We conclude that AtpΘ can associate with membranes despite the absence of a predicted membrane-spanning region.

**Figure 3.**
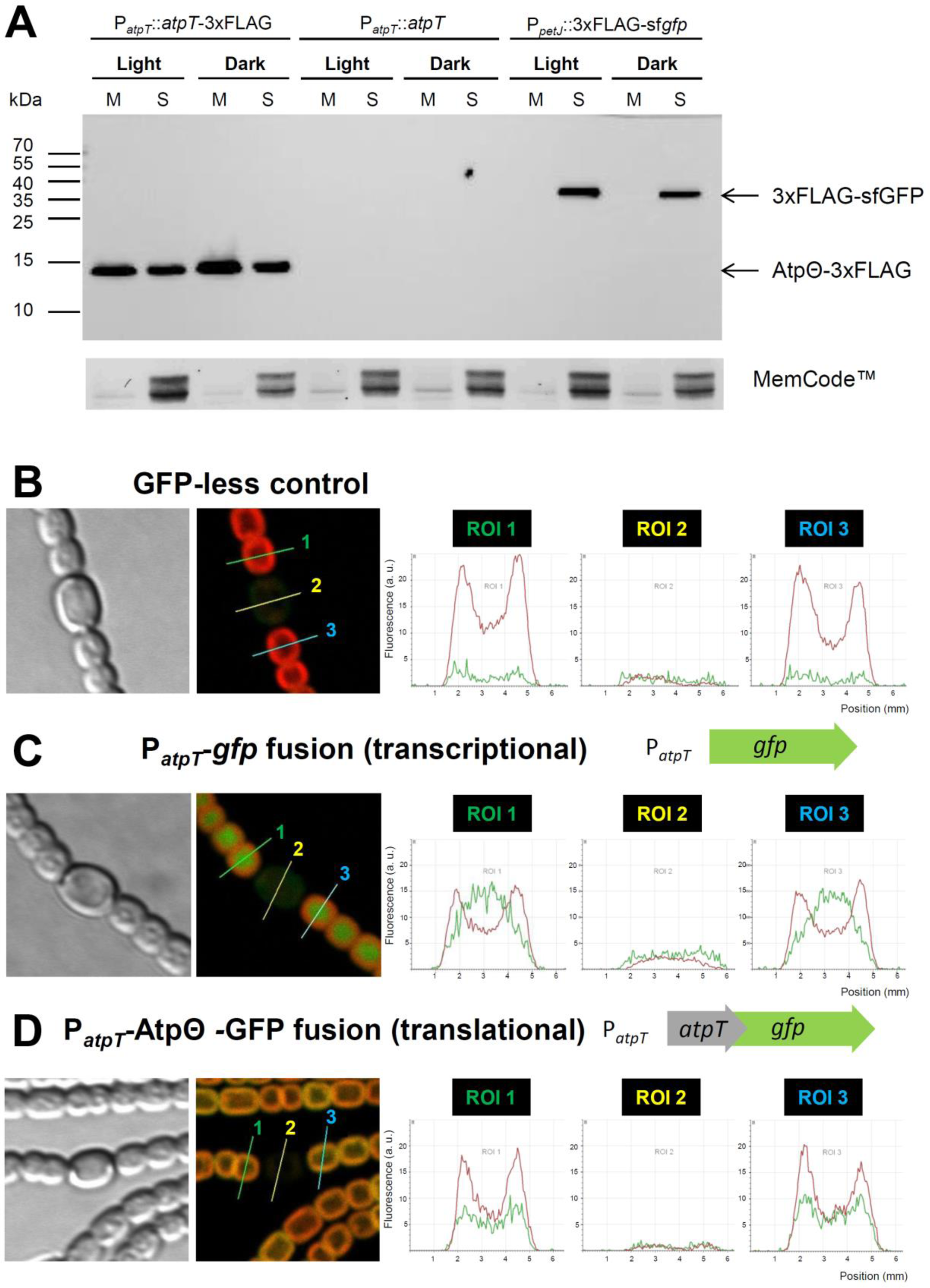
Intracellular localization of AtpΘ. **(A)** Localization of FLAG fusion proteins by separation of the membrane fraction from soluble proteins. Samples for protein extraction were taken from light- and dark (12 h incubation)-grown cultures of *Synechocystis* 6803 mutant strains: P_*atpT*_::*atpT*-3xFLAG and P_*petJ*_::3xFLAG-sf*gfp* expressed the recombinant AtpΘ and Gfp-FLAG fusion proteins, respectively, whereas P_*atpT*_::*atpT* was used as a negative control. Proteins (10 μg) were separated on a 15% (w/v) glycine SDS polyacrylamide gel and transferred to a nitrocellulose membrane, which was probed with specific ANTI-FLAG® M2-Peroxidase (HRP) antibody. MemCode™ Reversible Protein staining was used to check for equal protein loading. M, membrane fraction, S, soluble fraction. **(B) to (D)** Fluorescence-based analysis of the localization of AtpΘ in *Nostoc* 7120 bearing different fusions to GFP. **(B)** GFP-less control. **(C)** Transcriptional fusion: The *gfp* gene was placed under the control of the *atpT* promoter (construct pSAM342, **Table S2**). **(D)** Translational fusion: The *gfp* gene was fused to the *atpT* coding region and placed under the control of the *atpT* promoter (pSAM344, **Table S2**). In panels **(B)** to **(D)**, first, light transmission microscopy is shown, followed by fluorescence in the GFP channel merged with chlorophyll autofluorescence. The following three diagrams show the fluorescence intensities in cross sections region of interest (ROI) 1 to ROI 3 in three consecutive single cells, two vegetative cells and one heterocyst in the middle. GFP fluorescence is depicted in green, and chlorophyll autofluorescence is depicted in red.

### Fusions to AtpΘ target GFP to the cyanobacterial thylakoid membrane

According to the fractionation analysis in *Synechocystis* 6803, AtpΘ localizes to soluble and membrane-enriched protein fractions, but it remained unclear if only to the thylakoids, the cellular inner or outer membrane, or several of them. To obtain insight into the possible subcellular localization of AtpΘ, we chose *Nostoc* 7120 because of its much larger cells than *Synechocystis* 6803. TblastN analyses indicated the presence of a single possible *atpT* homolog in a chromosomal region to which a transcriptional start site was previously assigned at position 2982087r^30^. Northern hybridization showed a transcript originating from this region (**Figure 2F**), consistent with the length of 316 nt predicted for this gene from the TSS to the end of a Rho-independent terminator^31^. The corresponding gene was classified as protein-coding^31^ based on analysis by the RNAcode algorithm^32^. Upon shifting the cultures to darkness, this mRNA was rapidly induced (**Figure 2F**), similar to the regulation of the *atpT* gene in *Synechocystis* 6803 and three other cyanobacteria. Next, two constructs were prepared: pSAM342 harboring the *atpT* promoter, the corresponding 5’UTR plus the coding sequence for the green fluorescent protein (GFP) and pSAM344 harboring the *atpT* promoter, the 5’UTR, and the *atpT* coding region translationally fused to GFP (**Table S2**).

These constructs were introduced into plasmid α of *Nostoc* 7120 by homologous recombination. Confocal microscopy revealed GFP fluorescence in the recombinant strains obtained but not in a strain bearing a *gfp*-less control construct (**Figure 3B**).

However, we noticed a distinct difference in the intracellular localization of the signal. The fluorescence of the cells expressing the transcriptional fusion from construct pSAM342 appeared distributed throughout the cytoplasm, i.e., typical for a soluble protein such as GFP (**Figure 3C**). In contrast, the fluorescence of the translational fusion pSAM344 was localized differently and appeared spatially associated with the thylakoid membrane system, indicated by the overlap between chlorophyll and GFP fluorescence signals (**Figure 3D**). We conclude that translational fusions between *atpT* and *gfp* were translated well and that the AtpΘ sequence was competent to direct GFP to the thylakoid membrane. This localization is consistent with the association of soluble AtpΘ with a thylakoid membrane-bound complex.

Interestingly, for both constructs, the signal was very low in those cells that exhibited no chlorophyll fluorescence (compare **Figure 3B** with **Figures 3C** and **3D**). These cells were heterocysts specialized for nitrogen fixation, the assimilation of nitrogen from dinitrogen gas, N2, through the enzyme nitrogenase. This result provided evidence that the *atpT* promoter was switched off cell type-specifically in heterocysts.

### ATP synthase subunits become enriched in coimmunoprecipitation experiments

To identify the function of AtpΘ, protein coimmunoprecipitation assays followed by mass spectrometry were conducted with protein extracts from *Synechocystis* 6803 cells expressing FLAG-tagged AtpΘ under the control of its native promoter (strain P*atpT::atpT*-3xFLAG). As controls, a strain expressing untagged AtpΘ under control of the native promoter (strain P*atpT::atpT*) and a strain expressing FLAG-tagged sfGFP controlled by the copper-regulated P_*petJ*_ promoter (strain P_*petJ*_::3xFLAG-sf*gfp*) were used.

The evaluation of pull-down experiments via mass spectrometry showed that 34 proteins, including eight subunits of FoF1 ATP synthase, were enriched with a log_2_FC >3.5 among the proteins copurified with FLAG-tagged AtpΘ compared to at least one of the two controls (**Table S4**). These results pointed at a possible interaction between AtpΘ and one or several subunits of the FoF1 ATP synthase complex.

Two experiments were performed to verify this possibility. We performed a second immunoprecipitation assay comparing FLAG-tagged AtpΘ and FLAG-tagged sfGFP in three biological replicates each. The eluted samples were subjected to SDS-PAGE (**Figure S2**) and then analyzed using mass spectrometry. This analysis detected the same eight subunits of ATP synthase that were significantly enriched by coimmunoprecipitation with AtpΘ-3xFLAG (**Figure 4A**, marked in red), confirming the specific interaction between AtpΘ and the ATP synthase complex. Hence, in both analyses, the same 8 of the 9 known ATP synthase subunits were identified (**Tables S4** and **S5**). The only missing subunit was subunit c, the small membrane-intrinsic subunit, which appears to be difficult to detect by mass spectrometry. A small number of additional proteins significantly enriched by coimmunoprecipitation with AtpΘ-3xFLAG included two subunits of NAD(P)H-quinone oxidoreductase (subunit I and subunit O), two proteins of the CmpABCD transporter (CmpC and CmpA), and the bicarbonate transporter SbtA (**Figure 4B**), pointing at possible higher-order structures or additional binding partners of AtpΘ. The hierarchical clustering of AtpΘ-3xFLAG-enriched proteins labeled in **Figures 4A** and **4B**, as well as 3xFLAG-sfGFP is shown in **Figure 4C**. The resulting heat map further helped to visualize the enrichment of each protein, and the blank region under the 3xFLAG-sfGFP cluster indicates that no such proteins were identified. The complete dataset of the two independent coimmunoprecipitation assays can be obtained from the PRIDE partner repository (dataset identifiers PXD020126 and PXD024905).

**Figure 4.**
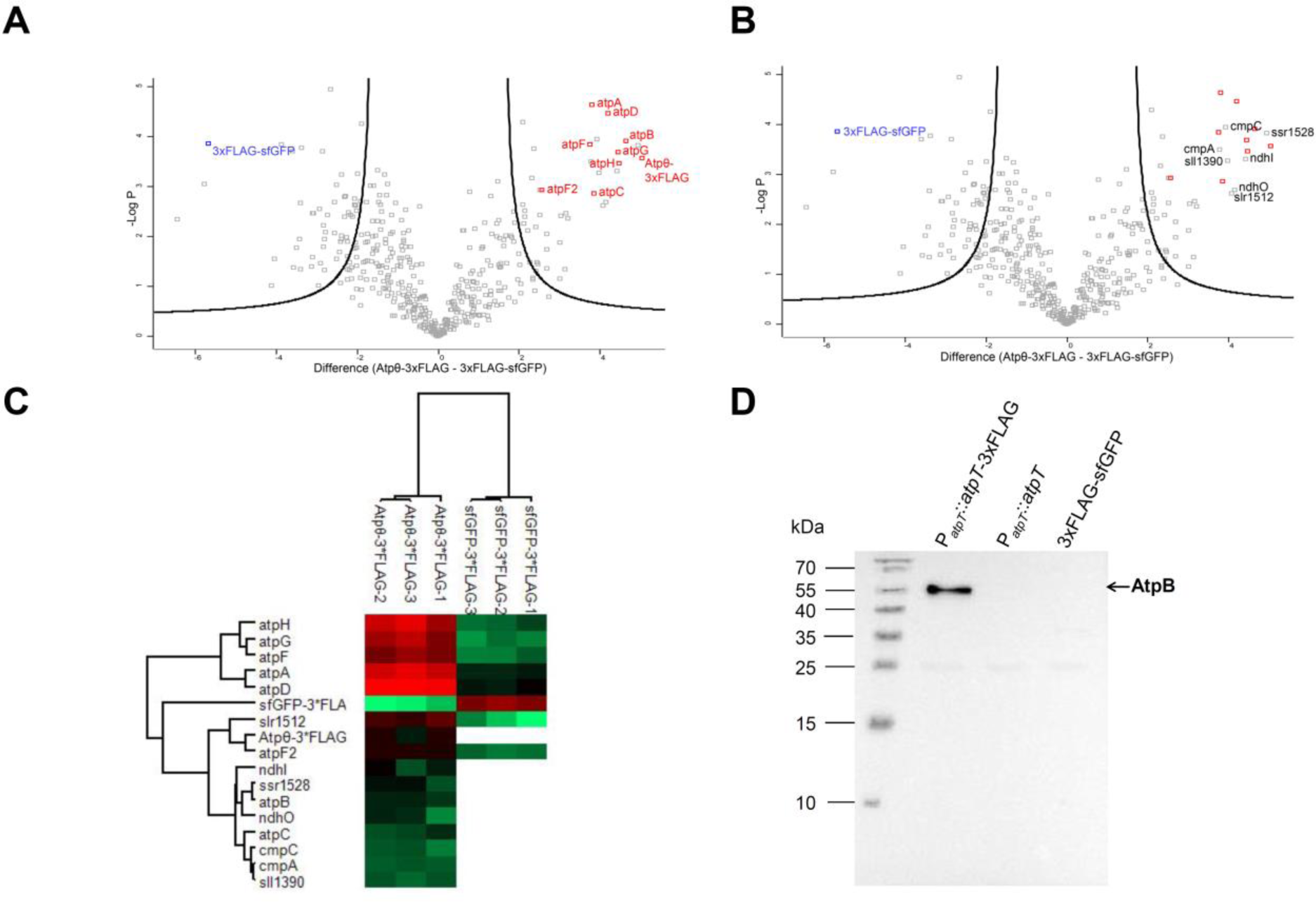
Copurification of AtpΘ and the ATP synthase complex verified by mass spectrometry and immunoblot analysis. **(A)** Volcano plot generated based on a two-sample *t*-test of enriched proteins using a false discovery rate (FDR) of 0.01 and a coefficient for variance minimization *s*_0_ ^48^ of 2. Atpθ-3xFLAG and the identified subunits of F_0_F_1_ ATP synthase are marked in red, while 3xFLAG-GFP is marked in blue. Subunit b’ (AtpF2) was added manually to the plot since this subunit was detected in only 2 out of 3 replicates. **(B)** The same volcano plot shown in **(A)** labeled with non-ATP synthase proteins. **(C)** Clustering heat map of the Atpθ-3xFLAG-enriched proteins marked in volcano plots (**A** and **B**). The log_2_ transformed NSAF intensities are indicated by different colors as indicated below. Undetected proteins in the 3xFLAG-GFP-enriched samples were left blank. 3xFLAG-GFP was detected in the Atpθ-3xFLAG group due to their common 3xFLAG tag. **(D)** Probing the elution fractions of P_*atpT*_::*atpT*-3xFLAG, P_*atpT*_::*atpT* and P_*petJ*_::3xFLAG-sf*gfp* with anti-AtpB serum.

As a further control experiment, we tested the enrichment of AtpB (subunit beta of ATP synthase) in an eluate from the pull-down experiment with FLAG-tagged AtpΘ by Western blotting. AtpB was clearly detected in this eluate but not in the eluate from the immunoprecipitation of 3xFLAG-sfGFP or a mock experiment with untagged AtpΘ (**Figure 4D**). Collectively, these results supported an interaction between AtpΘ and subunit(s) of the ATP synthase complex. Moreover, this interaction would explain the association of AtpΘ with thylakoid membranes as was observed in **Figure 3**.

### Impact of AtpΘ on ATP synthase activity

The results in **Figure 3** showed that AtpΘ associates with thylakoid membranes and the results in **Figure 4** that it is the ATP synthase complex it is interacting with. To test its functional impact, the *atpT* gene was replaced by a chloramphenicol resistance cassette and biochemical measurements of ATPase activity were performed. Membrane fractions were isolated from both the wild type and the fully segregated *atpT* knockout strain (**Figure S1D),** which had been kept in continuous light or dark, and their ATP hydrolysis activities were analyzed. The results (**Figure 5A**) showed that the membrane fraction of wild-type *Synechocystis* 6803 grown in the light had a significantly higher ATPase activity than the membrane fraction isolated after 24 h of darkness incubation. In contrast, the membrane preparation from the knockout strain without AtpΘ showed no significant difference between the light- and dark-incubated conditions. These results suggested an *in vivo* inhibitory effect of AtpΘ on ATPase activity under darkness.

**Figure 5.**
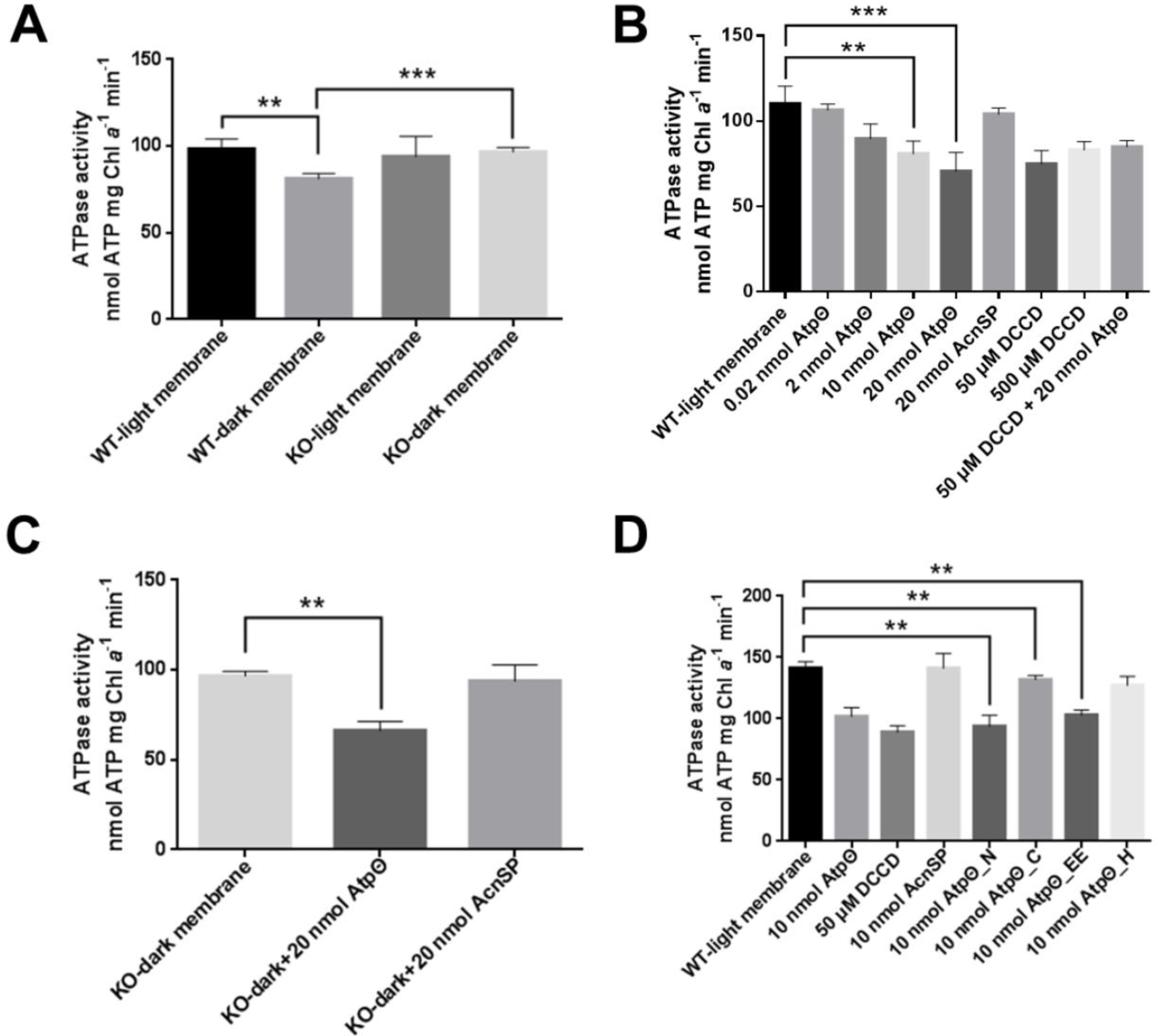
ATPase activities in membrane fractions. **(A)** ATPase activity of the membrane fraction of wild type and *atpT* knockout *Synechocystis* 6803 cells growing under continuous light or after 24 hours of darkness incubation. **(B)** ATPase activity of the membrane fraction of wild-type *Synechocystis* 6803 supplemented with different synthetic peptides or chemicals. DCCD was used as a positive control for ATPase activity inhibition, while the synthetic AcnSP peptide was used as a negative control. **(D)** ATPase activity of the membrane fraction isolated from *atpT* knockout *Synechocystis* 6803 cells after 24 hours of darkness incubation supplemented with either synthetic AtpΘ or AcnSP peptide. **(D)** ATPase inhibitory effects of AtpΘ peptides with truncated or modified sequences (**Figure S4**). The differences between groups were tested using GraphPad software as described in the methods section. Significance was established at *P* < 0.05 = ** and *P* < 0.01 = ***.

Similar findings were observed in a second cyanobacterium, *Thermosynechococcus elongatus* BP-1, where the ATPase activities of the membrane samples prepared from light-cultivated cells were significantly higher than the ATPase activities of the membrane samples from dark-incubated cells (**Figure S3**). Thus, the predicted AtpΘ homolog of *Thermosynechococcus elongatus* BP-1 could function similarly to AtpΘ of *Synechocystis* 6803.

To further characterize the potential inhibitory effect of AtpΘ, the ATP hydrolysis activity of the membrane fraction from wild-type *Synechocystis* 6803 cells was measured in the presence of different amounts of an AtpΘ synthetic peptide (**Figure 5B**). The synthetic peptide AcnSP^33^, which is of a length similar to AtpΘ and was synthesized by the same company, was used as negative control. In parallel, the well-established FoF1 ATP synthase inhibitor DCCD served as positive control. As shown in **Figure 5B**, AtpΘ reduced ATPase activity in a dose-dependent manner, and the inhibitory effect was saturated at 20 nmol AtpΘ, whereas the AcnSP peptide showed no effect on ATPase activity. High amounts of DCCD inhibited ATPase activity at a level similar to the AtpΘ peptide. Finally, the combination of DCCD and AtpΘ peptide yielded an ATPase inhibition similar to their separate addition. These results indicated that AtpΘ is a strong inhibitor of the ATP hydrolysis activity of FoF1 ATP synthase, comparable to DCCD. The remaining 60% ATP hydrolysis activity of the membrane preparations probably resulted from other ATP hydrolases, such as H^+^-translocating P-type ATPases that are resistant to DCCD or PilT1 and PilB1 proteins providing energy for the type IV pili system^34, 35^, or because either AtpΘ or DCCD cannot fully inhibit ATPase activity. Then, to further confirm whether the difference between wild-type and knockout cells observed in **Figure 5A** was due to the lack of AtpΘ, 20 nmol of AtpΘ or AcnSP peptides was supplemented to the membrane isolated from the dark-incubated knockout strain. The results showed that supplementation with AtpΘ could significantly inhibit the ATPase activity of the membrane, while AcnSP showed no such effects (**Figure 5C**), further confirming the inhibitory role of AtpΘ.

To identify the minimal inhibitory sequence of AtpΘ and to study the effects of specific amino acids on the ATPase inhibitory effect of AtpΘ, four mutant AtpΘ peptides were designed and synthesized (**Figure S4**). The inhibitory effects of these peptides on ATPase activity were tested and compared to the inhibitory effects of the original AtpΘ peptide (**Figure 5D**). Interestingly, the N-terminal part of AtpΘ, which corresponds to the predicted alpha-helical part of this protein (peptide AtpΘ_N in **Figure 5D** and **Figure S4**), exhibited an inhibitory effect similar to that of? the entire peptide. In contrast, the central part of AtpΘ (AtpΘ_C in **Figure 5D** and **Figure S4**) showed weaker inhibitory effects. Introduction of two conserved amino acid substitutions (D26E and D27E) yielded AtpΘ_EE, as shown in **Figure 5D** and **Figure S4**. Consistent with the conservative replacement of two acidic residues by two others, a similar inhibitory effect on ATP hydrolysis activity was observed as for the native AtpΘ protein. In contrast, the introduction of a single histidine residue at this position (E27H; AtpΘ_H in **Figure 5D** and **Figure S4**) led to an almost complete loss of the inhibitory effect, indicating that the negative charge at this position is important for the inhibitory activity of the full-length AtpΘ peptide. The 3D structure modeling of AtpΘ using PEP-FOLD3^36^ and analysis using the HELIQUEST web server^37^ predicted an N-terminal amphipathic alpha helix and a C-terminal random structure, which was also observed in the predicted structures of representative AtpΘ homologs from cyanobacteria strains with one, two or three putative homologs (**Figure S5**). These results suggest that the N-terminal alpha helix of AtpΘ is a conserved structural element that is together with the cluster of centrally located, negatively charged amino acids responsible for the inhibition of ATPase activity.

### F_0_F_1_ ATP synthase purification and the effect of AtpΘ

To rule out the effects of membrane proteins other than ATP synthase, ATP synthase was purified from *Synechocystis* 6803 cells by fusing a 3xFLAG tag to the C-terminus of AtpB. The purified protein complex was first characterized by SDS-PAGE, showing good purity and distribution of different subunits (**Figure 6A**), and then probed using anti-FLAG and anti-AtpB antisera (**Figures 6B** and **6C**, respectively), confirming the presence of both 3xFLAG and AtpB. The ATPase activity of the purified ATP synthase was then measured directly or in the presence of different inhibitors (**Figure 6D**). ATP hydrolysis activity was detected using the purified protein complex. Compared with the purified complex, the addition of AcnSP peptide yielded no significant changes, whereas the addition of AtpΘ peptide or DCCD significantly decreased the ATPase activity. The inhibitory effect of AtpΘ appeared stronger than the inhibitory effect of DCCD, and the combination of both showed no additive effects. These results further confirmed the inhibitory effect of the AtpΘ peptide on the hydrolytic activity of ATP synthase.

**Figure 6.**
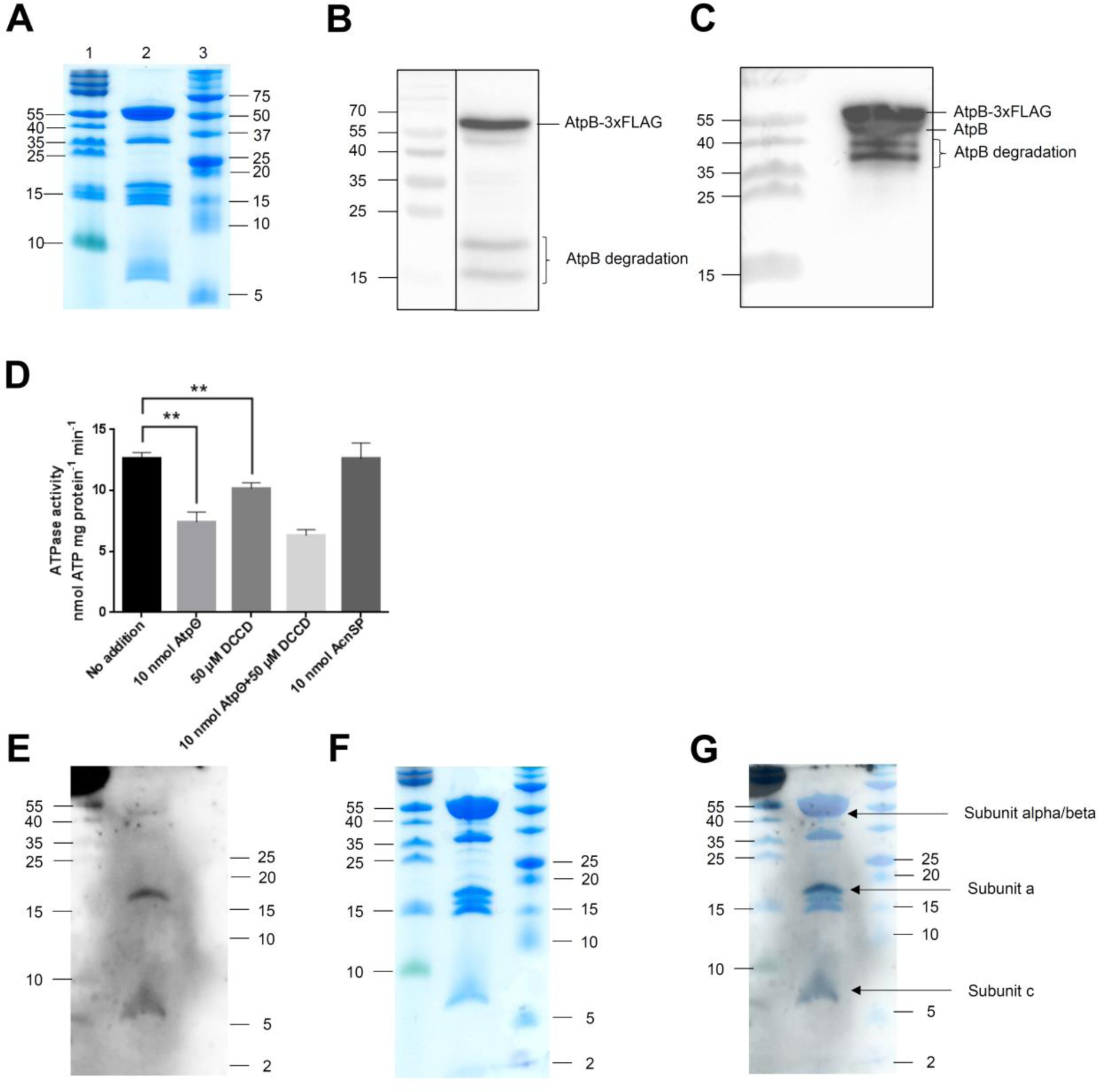
Purification of the F_0_F_1_ ATP synthase and its interaction with AtpΘ. **(A)** Tricine SDS-PAGE displaying the purity of 10 μg *Synechocystis* 6803 F_0_F_1_ ATP synthase (lane 2) isolated with AtpB-3xFLAG and gel filtration chromatography. PageRuler™ Prestained Protein Ladder (lane 1; 10 to 180 kDa) and Precision Plus Protein™ Dual Xtra Prestained Protein Standard (lane 3; 2 to 250 kDa) were used as molecular mass markers. **(B)** Western blot analysis of 10 μg purified *Synechocystis* 6803 F_0_F_1_ ATP synthase probed with specific ANTI-FLAG® M2-Peroxidase (HRP) antibody. **(C)** Western blot analysis of 10 μg purified *Synechocystis* 6803 F_0_F_1_ ATP synthase probed with anti-AtpB serum. The doublet for AtpB consists of the native and 3xFLAG-tagged forms of the protein. Bands in panels b and c likely resulting from degradation are labeled. **(D)** Measurement of the ATP hydrolysis activity of purified F_0_F_1_ ATP synthase supplemented with different synthetic peptides or chemicals. DCCD was used as a positive control for ATPase activity inhibition, while the synthetic AcnSP peptide was used as a negative control. The differences between groups were tested using a paired *t*-test (**Table S10**) using GraphPad software. Significance was established at **, *P* < 0.05. **(E)** Far Western blot signal detecting the interaction partners of synthetic AtpΘ peptide from purified *Synechocystis* 6803 F_0_F_1_ ATP synthase. **(F)** Coomassie blue staining of the Tricine-SDS gel after blotting. **(G)** The immunoblot signal (**E**) was merged with the stained gel (**F**) to determine the interacting subunits. The subunits suspected to interact with Atpθ are labeled. PageRuler™ Prestained Protein Ladder (10–170 kDa, Fermentas; left) and Protein™ DualXtra (2-250 kDa, Bio-Rad; right) were used as molecular mass markers. In total, 15 μg purified ATP synthase was loaded on the gel.

### Far Western blot identifies interaction partners of AtpΘ from purified ATP synthase

To gain further insight into the interaction between AtpΘ and the ATP synthase complex, a far Western blot approach^38^ was applied. In this approach, proteins were renatured after blotting onto a membrane and served as baits. The membrane was then incubated with the synthetic AtpΘ peptide, followed by anti-AtpΘ serum and anti-rabbit IgG antiserum. As shown in **Figure 6E-G**, AtpΘ was enriched mainly at two positions, which were assigned as subunit a (*atpB*, Sll1322) and subunit c (*atpE*, Ssl2615), respectively, based on the comparison to the subunit distribution of *E. coli* F_0_F_1_ ATP synthase^39^. A very weak signal of approximately 50 kDa size was also observed, which should correspond to subunit alpha or subunit beta (**Figure 6G**). Although the 50 kDa signal was relatively weak, it was reproducibly observed (n = 3) and should therefore also be considered. As a negative control, a mock far Western blot was conducted in which TBST buffer was used instead of the synthetic AtpΘ peptide. In this setting, no signal was observed for the two replicates of ATP synthase purification, while the positive controls could be detected (**Figure S6**), suggesting that none of the signals observed in **Figures 6E and 6G** were due to unspecific interaction with any of the antisera used. These results further confirmed the specific interaction of the AtpΘ peptide with distinct subunits of ATP synthase.

## Discussion

F_0_F_1_-type ATP synthases produce ATP via chemiosmotic coupling to a proton gradient. However, while ATP synthases preferentially catalyze ATP formation, a weaker or temporarily missing proton gradient can stimulate the reverse reaction, pumping protons while hydrolyzing ATP. In the mitochondria of yeast and mammals, small inhibitory peptides can prevent ATP synthase from running backwards, hence avoiding wasteful ATP hydrolysis. In plant chloroplasts, in which the proton gradient is generated by light-driven photosynthetic electron transport, a redox-controlled mechanism switches ATP synthase activity off at night when photosynthesis does not take place^1^. In this respect, cyanobacteria present an interesting case because photosynthetic ATP synthesis is light-driven, as in chloroplasts, but conditions can easily be envisioned where inhibition of ATP hydrolysis is warranted. Such conditions can be low light, darkness and others that would affect the strength of the proton gradient. Cellular ATP demand is likely considerably lower under certain conditions, e.g., during the night, but in contrast to chloroplasts, residual activity should be maintained in cyanobacteria to allow respiration to proceed with ATP synthesis, which uses the same ATP synthase that is also used for photosynthesis in the thylakoid membrane system.

Several mechanisms have been identified for regulating ATP synthase activity in cyanobacteria (**Figure 7**). A common regulatory mechanism, the ADP-mediated inhibition of the F1 part, has been reported for cyanobacterial ATPase^20^. Although the γ subunit of cyanobacteria is not redox-sensitive compared to the chloroplast F_0_F_1_ ATP synthase subunit γ, the ADP-mediated inhibition of ATPase was assigned to this subunit^20^. Another well-characterized mechanism is the inhibition of the rotation of bacterial F_0_F_1_ ATP synthase via the ε subunit, called ε inhibition^40^. ε inhibition in cyanobacteria was reported to be ATP-independent, different from other bacteria, and was related to the distinct γ subunit of cyanobacteria as well^41^. In addition, both the γ and ε subunits of cyanobacterial F_0_F_1_ ATP synthase were reported to be important for the dark acclimation of cyanobacteria^21^. However, many aspects of the regulation of cyanobacterial F_0_F_1_ ATP synthase have remained unknown.

**Figure 7.**
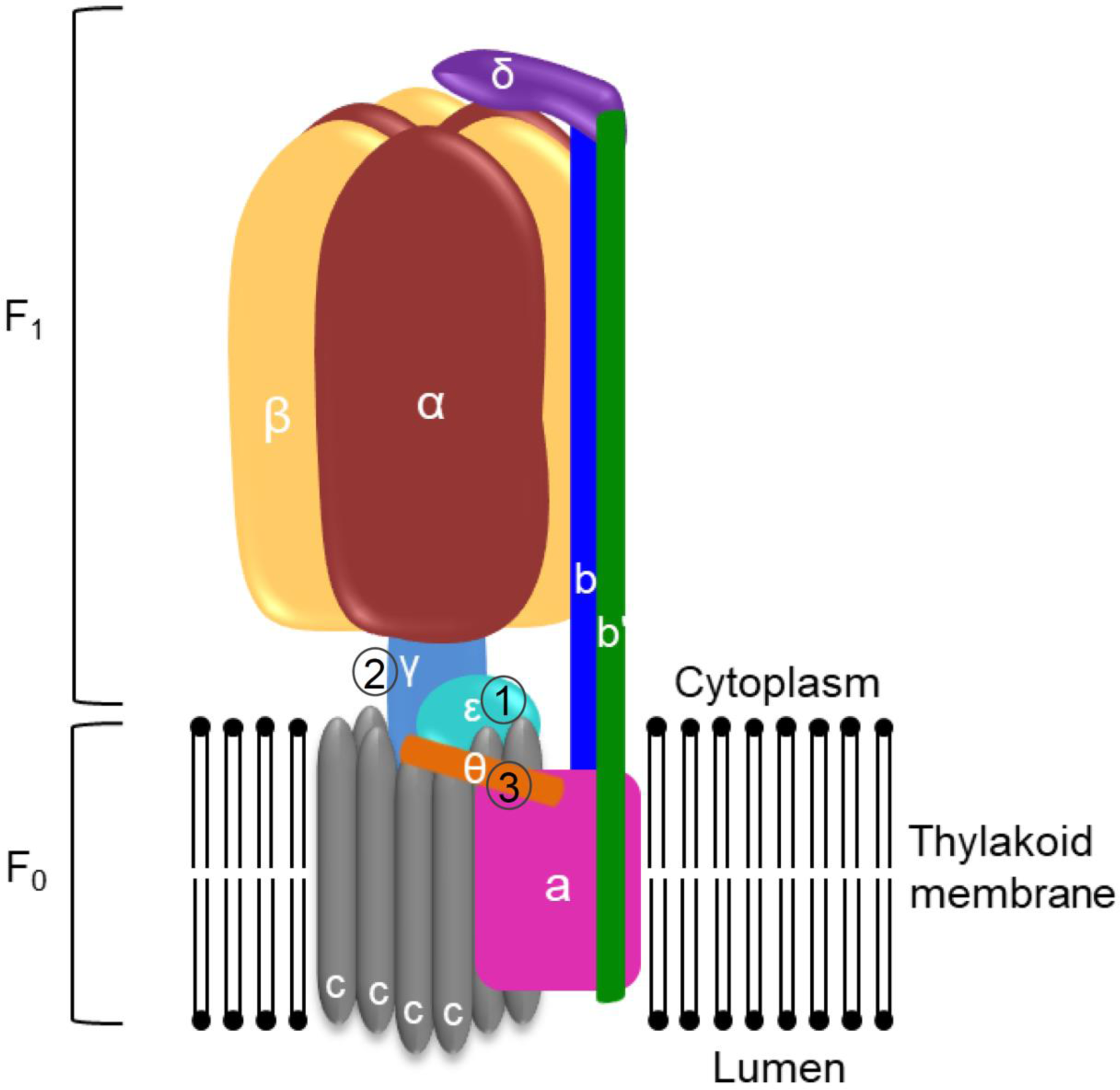
Regulatory and inhibitory mechanisms impacting hydrolysis of ATP by ATP synthases in cyanobacteria. (1) ε–inhibition^21^, a mechanism in which the ε subunit inhibits the rotation of the γ subunit via a conformational change^41^; (2) ADP inhibition, the most conserved mechanism to block ATPase activity (reviewed by Lapashina and Feniouk^49^) via a segment inserted in the γ subunit^20^, or (3) by AtpΘ. In this suggested mechanism, H^+^ ions pass through the c ring and drive its rotation, and then the upper part of the complex rotates together in the non-inhibited state. AtpΘ may block the rotation of the c ring via interaction with subunit a. The computationally predicted structure of AtpΘ has tentatively been drawn next to subunits a and c, but the exact topography and mode of interaction with these membrane-embedded subunits is a topic of future research.

In the present study, we suggest that the small protein AtpΘ represents a functional analog in cyanobacteria of the small inhibitory peptides that arrest ATP synthase from running backwards in mitochondria. This hypothesis is supported by several lines of evidence. First, the direct interaction of AtpΘ with ATP-synthase subunits has been shown in different protein/protein interaction studies, where it showed the strongest binding toward subunit a (*atpB*, Sll1322) and subunit c (*atpE*, Ssl2615) (**Figure 6G**). Second, AtpΘ supplementation had a specific, dose-dependent negative impact on ATP hydrolysis activity in isolated membrane fractions or using the purified ATP synthase complex. The extent of this inhibition was similar to the inhibition exerted by specific ATP synthase inhibitors, consistent with its role in preventing the reverse reaction, i.e., the wasting of ATP via hydrolysis when the proton gradient is weakened. The addition of synthetic Atpθ peptide to the membrane samples prepared from *Synechocystis* 6803 cultured in the presence of light yielded a maximum 35% inhibition of ATPase activity (**Figure 5B**). A slightly higher inhibitory effect of 40% was achieved if Atpθ peptide was added to the preparations of purified ATP synthase (**Figure 6D**). These findings indicate that the Atpθ peptide may inhibit up to 40% of the ATPase activity in ATP synthase. Third, investigations of wild type and *atpT* mutant strains are consistent with the *in vitro* ATP synthase activity tests. While the membrane sample prepared from *Synechocystis* 6803 cultured under continuous light showed a similar ATPase activity as previously reported^21^, the membrane samples prepared from dark-incubated *Synechocystis* 6803 with maximum *atpT* expression showed an ATPase activity of approximately 85% (**Figure 5A**), and this difference was not observed in the *atpT* knock-out mutant of *Synechocystis* 6803, indicating that the Atpθ protein is required *in vivo* to decrease ATPase activity during dark incubation. Fourth, the expression data suggest that the AtpΘ action is controlled mostly via the regulation of its expression because the protein seems to have a low stability (**Figure 2D**). Its expression is particularly stimulated under conditions that could weaken the transmembrane proton gradient, such as darkness, or, in our experiments, by the addition of the uncoupler CCCP or the electron chain inhibitor DBMIB (**Figure 2B**). While CCCP is a well-established protonophore, DBMIB is better known as an inhibitor of photosynthetic electron transfer. However, DBMIB affects the cytochrome *b6f* complex^27^, which is shared by photosynthetic and respiratory electron transfer that operates in the same membrane system in cyanobacteria. In contrast, the presence of AtpΘ might be futile when ATP synthase runs at high speed, such as under high-light conditions driven by an efficient photosynthetic light reaction or in the presence of high rates of respiration. Consistently, *atpT* dark induction could largely be prevented by the addition of glucose (**Figure 2A**), likely due to the stimulation of respiration-dependent ATP synthesis in the presence of glucose. Furthermore, such a scenario of high respiration exists in heterocysts, which have no photosystem II but exhibit substantial ATP production to meet the high demand of nitrogen-fixing nitrogenase, linked to high rates of respiration consuming the O_2_ inside the heterocysts. Indeed, we observed that the expression of the *atpT* promoter was shut down in heterocysts of *Nostoc* 7120 (**Figure 3C** and **3D**). Finally, the cross-phylum importance of the AtpΘ-mediated prevention of the backward reaction of ATP synthase is supported by its ubiquitous occurrence throughout the cyanobacterial phylum and our finding that *atpT* expression was stimulated under conditions leading to lowered thylakoid proton gradients in several divergent species of cyanobacteria (**Figure 2E** and **2F**). These results make it very likely that the conclusions obtained with the model *Synechocystis* 6803 can also be generalized for other cyanobacteria.

Collectively, these data provide evidence that the small protein AtpΘ acts as an ATP hydrolysis inhibitor of cyanobacterial ATP synthase. This role of AtpΘ represents an interesting analogy to the ATP synthase regulator IF1 in the mitochondria of eukaryotes. However, unlike IF1, which binds the catalytic interface between the α and β subunits^42^, the main potential interaction partners of AtpΘ, subunits a and c, belong to the F0 part of ATP synthase, which resides within the thylakoid membrane (**Figure 7**). The predicted amphipathic character of the N-terminal helix in AtpΘ homologs (**Figure S5C**) may support the interaction with membrane proteins such as subunits a and c but may facilitate also additional interactions with subunits in the soluble phase. Therefore, the binding of AtpΘ to the two topographically close subunits a and c and its amphipathic character point to a possibly divergent mechanism of AtpΘ function by hindering the rotation of the ATP synthase complex through direct binding.

## Acknowledgments

This study was funded by the German Research Foundation (DFG) priority program SPP2002 “Small Proteins in Prokaryotes, an Unexplored World” (grant HE 2544/12-1 to WRH, grant HA 2002/22-1 to MH and grant BE 3869/5-1 to DöB), by grant PID2019-105526GB-I00/AEI/10.13039/501100011033 (AEI/FEDER, UE) to AMMP, and by a China Scholarship Council grant to K.S. We thank the ZBSA proteomics staff, especially Verónica I. Dumit, for support, Gen Enomoto, for an aliquot of *Thermosynechococcus elongatus* BP1, Mai Watanabe for an aliquot of *Gloeobacter violaceus* PCC 7421 and Claudia Steglich for a culture of *Prochlorococcus* sp. MED4 (all University of Freiburg).

## Author contributions

KS and DeB carried out the molecular-genetic and biochemical analyses in *Synechocystis* 6803, and AMMP performed all experiments in *Nostoc* 7120. SM and DöB performed proteomics analyses. MH provided scientific input for improving the experimental design and physiological interpretation. WRH designed the study, and all authors analyzed the data. KS and WRH drafted the manuscript. All authors read and approved the final manuscript.

## STAR Methods

### RESOURCE AVAILABILITY

#### Lead contact

Further information and requests for resources and reagents should be directed to and will be fulfilled by the lead contact, Wolfgang R. Hess.

#### Materials availability

N/A

#### Data and code availability

- Mass spectrometry raw data have been deposited at the ProteomeXchange Consortium (http://proteomecentral.proteomexchange.org) and are publicly available as of the date of publication. Accession numbers are listed in the key resources table.
- This paper does not report original code.
- Any additional information required to reanalyze the data reported in this paper is available from the lead contact upon request.

### EXPERIMENTAL MODEL AND SUBJECT DETAILS

#### Cultivation of cyanobacteria

Wild-type *Synechocystis* sp. PCC 6803 PCC-M and mutant strains were cultured photoautotrophically in TES-buffered (20 mM, pH 8.0) BG11 medium^50^ with gentle agitation or on agar-solidified (1.5% Kobe I agar) plates under constant illumination with white light of approximately 40 µmol photons m^-2^ s^-1^ at 30°C and supplemented with appropriate antibiotics (5 µg/mL gentamicin, 10 µg/mL kanamycin, and 3 µg/mL chloramphenicol). For incubation in darkness, flasks were wrapped with aluminum foil. CuSO_4_ (2 μM) was used to induce the expression of the Cu^2+^-responsive *petE* promoter^51^, while the *petJ* promoter was induced by removing Cu^2+^ from the medium through centrifugation and resuspension. For high-density cultivation used for ATP synthase purification, *Synechocystis* 6803 overexpressing P*petE*-*atpB*-3xFLAG was cultured in the cell-DEG system as reported previously^52^ using freshwater medium^53^ with the following modifications: Na2EDTA and CuSO_4_ were not included in the medium, and 10 µg/mL kanamycin or 5 µg/mL gentamicin was added.

Cultures of *Nostoc* 7120 were bubbled with an air/CO_2_ mixture (1% v/v) and grown photoautotrophically at 30°C in BG11 medium^50^. Darkness was implemented on air-CO_2_-bubbled cultures by covering with aluminum foil plus black velvet. The thermophilic cyanobacterium *Thermosynechococcus elongatus* BP-1 was cultured in BG11 medium under continuous illumination with 30 µmol photons m^-2^ s^-1^ white light (Master LED tube Universal 1200 mm UO 16 W830 T8; Philips) at 45°C. *Gloeobacter violaceus* PCC 7421 was cultivated photoautotrophically in Allen’s medium^54^ in Erlenmeyer flasks under continuous white light (4 μmol photons m^−2^ s^−1^) at 20°C with shaking. *Prochlorococcus* MED4 cells were grown at 22°C in AMP1 medium^55^ under 30 µmol photons m^-2^ s^-1^ continuous white cool light and harvested in an exponential growth phase.

### METHOD DETAILS

#### Construction of mutant cyanobacterial strains

To delete the *atpT* gene from *Synechocystis* 6803 (genome position 3274499 to 3274645, reverse strand), the flanking regions of *atpT* were amplified by primer pairs AtpTKO-up-F/AtpTCmKO-up-R and AtpTCmKO-down-F/AtpTKO-down-R, and the resulting fragments were fused with a chloramphenicol resistance cassette and a pUC19 backbone amplified with primer pairs AtpTKO-vec-F/AtpTKO-vec-R and CmR-F/CmR-R using AQUA cloning^56^. The resulting plasmid, pUC-atpTKO-CmR, was then transferred into wild-type *Synechocystis* 6803 by natural transformation. The transformants were selected on BG11 agar plates supplemented with chloramphenicol. Complete segregation was achieved after several rounds of selection.

The construction of overexpression strains P_*atpT*_::*atpT*, P_*atpT*_::*atpT*-3xFLAG, and P_*petJ*_::3xFLAG-sf*gfp* was described previously ^24^. P*petE*::*atpB*-3xFLAG, a strain overexpressing the FLAG-tagged subunit AtpB, was constructed using primer pairs pUC19-XbaI_PpetE_fw/atpB::PpetE_rev, PpetE::atpB_fw/3xFlag_atpB_rev and atpB_3xFlag_fw/3xFlag_PstI-pUC19_rev. The primers used for mutant construction are listed in **Table S3**.

Construction of *Nostoc* 7120 strains. The strain carrying plasmid pCSEL24 (*gfp*-less control) was constructed previously^57^. To construct pSAM342 (transcriptional fusion of coordinates 2982431 to 2982070 from *Nostoc* 7120, reverse strand), a PCR fragment was amplified with oligonucleotides 900+901, *Cla*I-*Xho*I-digested and cloned into *Cla*I-*Xho*I-digested pSAM270^58^. To generate pSAM344 (translational Atpθ-GFP fusion, coordinates 2982431 to 2981902, reverse strand), a PCR fragment was amplified with oligonucleotides 900+902, *Cla*I-*EcoR*V-digested and cloned into *Cla*I-*EcoR*V-digested pSAM147^59^ in frame with the *gfpmut2* gene, rendering pSAM343. The *EcoR*I fragment from pSAM343, containing the fusion between the *atpT* promoter, the *atpT* gene and the *gfpmut2* gene, was cloned into *EcoR*I-digested pCSEL24, rendering pSAM344. All plasmids were transferred by conjugation followed by selection of streptomycin/spectinomycin (5 µg/mL each)-resistant colonies after integration in the alpha megaplasmid.

#### Computational sequence analyses

Homologs of the *atpT* gene were searched using the *Synechocystis* 6803 AtpΘ as query against the IMG^60^ and UniProt databases using blastP and against the NCBI database using both TblastN^61^ and blastP^62^ at a threshold E value ≤1e^-5^. Multiple sequence alignments were conducted using Jalview2^63^. Isoelectric points were predicted by the R package pIR^64^.

Phylogenetic analyses were conducted in MEGA X^44^ using the maximum likelihood algorithm based on 16S rRNA sequences extracted from the SILVA database^43^ and modified according to Klähn *et al*.^65^. The evolutionary distances were computed using the maximum composite likelihood method^66^.

#### Fluorescence microscopy

Images of *Nostoc* 7120 filaments growing on top of nitrogen-free solid media were taken five days after plating. The accumulation of GFP was analyzed and quantified using a Leica TCS SP2 confocal laser scanning microscope as previously described^67^.

#### Protein extraction and Western blots

*Synechocystis* 6803 cells for protein extraction were harvested by centrifugation (4,000 x g, 10 min, 4°C) and resuspended in PBS buffer (137 mM NaCl, 2.7 mM KCl, 10 mM Na_2_HPO_4_, 1.8 mM KH_2_PO_4_, pH 7.4) containing protease inhibitor cocktail. Cells were then disrupted mechanically in a Precellys homogenizer (Bertin Technologies). Glass beads and unbroken cells were removed by centrifugation at 1,000 g for 1 min at 4°C, and the total crude protein was obtained. Before loading, protein samples were boiled with 1x protein loading buffer at 95°C for 10 min or incubated at 50°C for 30 min supplemented with 2% SDS if the membrane fraction was included.

For Western blot analysis, proteins were separated either in 15% glycine-SDS gels or in 16%/6 M urea Tricine-SDS gels. PageRuler Prestained Protein Ladder (10– 170 kDa, Fermentas) or Precision Plus Protein DualXtra (2–250 kDa, Bio-Rad) was used as a molecular mass marker. The separated proteins were then transferred to nitrocellulose membranes (Hybond™-ECL, GE Healthcare) by semidry electroblotting. The blotted membrane was then blocked with 3% skimmed milk dissolved in TBST (20 mM Tris pH 7.6, 150 mM NaCl, 0.1% Tween-20) and incubated with primary antibody (1:500 dilution for anti-AtpΘ antiserum and 1:2,000 for anti-AtpB antibody) and secondary antibody (1:10,000 anti-rabbit antibody) sequentially. The anti-AtpΘ antiserum was generated by a commercial provider (Pineda Antikörper-Service). Signals were detected with ECL start Western blotting detection reagent (GE Healthcare) on a chemiluminescence imager system (Fusion SL, Vilber Lourmat).

#### Isolation of FLAG-tagged proteins and mass spectrometry analysis

For the pull-down assay, 800 mL of *Synechocystis* 6803 culture at an OD_750_ of approximately 1 was harvested by centrifugation at 4,000 g for 30 min at 4°C. Cell pellets were washed once with prechilled FLAG buffer (50 mM Hepes-NaOH pH 7.0, 5 mM MgCl2, 25 mM CaCl2, 150 mM NaCl, 10% glycerol, 0.1% Tween-20) and then resuspended in the same buffer supplemented with protease inhibitor cocktail. The cell suspension was disrupted with a Precellys homogenizer (Bertin Technologies, France). All subsequent steps were carried out at 4°C. Total cell extracts and glass beads were transferred to Bio-Spin® Disposable Chromatography Columns (Bio-Rad), which were put on centrifugation tubes (Sorvall Instruments). The glass beads were separated from cellular components by centrifugation (4,000 g, 5 min, 4°C). Membrane proteins were then solubilized by adding 2% *n*-dodecyl-beta-D-maltoside (β-DM), followed by dark incubation for 1 h at 4°C with gentle agitation. Nonsoluble components were removed by centrifugation (25,000 g, 30 min, 4°C), and the solubilized crude extract (sCE) was transferred to a new tube.

FLAG-tagged proteins were purified by column chromatography using ANTI-FLAG M2 affinity agarose gel or ANTI-FLAG M2 magnetic beads (both from Sigma-Aldrich) according to the manufacturer’s instructions. When ANTI-FLAG M2 affinity agarose gel was used, the sCE was loaded and passed over the column three times to improve binding. The column was washed with 5 x 2 mL of FLAG buffer, and the FLAG-tagged proteins were eluted by incubating the matrix with 1x protein loading buffer at 50°C for 30 min. The agarose was sedimented by centrifugation (16,000 g, 5 min, RT), and 20 μL of the resulting eluates were loaded on a glycine SDS-PAGE, which was subsequently stained with Coomassie. Then, each gel lane was cut into pieces, destained, desiccated and rehydrated in trypsin as previously described^68^. In gel-digest was incubated at 37 °C overnight and peptides were eluted with water by sonication for 15 min.

Samples were loaded on an EASY-nLC II system (Thermo Fisher Scientific) equipped with an in-house built 20 cm column (inner diameter 100 µm, outer diameter 360 µm) filled with ReproSil-Pur 120 C18-AQ reversed-phase material (3 µm particles, Dr. Maisch GmbH). Elution of peptides was achieved with a nonlinear 77 min gradient from 1 to 99% solvent B (0.1% (v/v) acetic acid in acetonitrile) with a flow rate of 300 nl/min and injected online into an LTQ Orbitrap XL (Thermo Fisher Scientific). The survey scan at a resolution of R=30,000 and 1 x 10^6^ automatic gain control target in the Orbitrap with activated lock mass correction was followed by selection of the five most abundant precursor ions for fragmentation. Single charged ions as well as ions without detected charge states were excluded from MS/MS analysis. Fragmented ions were dynamically excluded from fragmentation for 30 s. Database searches with Sorcerer-SEQUEST 4 (Sage-N Research, Milpitas, USA) were performed against a *Synechocystis* 6803 database downloaded from Uniprot (Proteome-ID UP000001425) on 11/12/20, which was supplemented with common laboratory contaminants and the sequences of AtpΘ-3xFLAG and GFP-3xFLAG. After adding reverse entries the final database contained 7,102 entries. Database searches were based on a strict trypsin digestion with two missed cleavages permitted. No fixed modifications were considered and oxidation of methionine was considered as variable modification. The mass tolerance for precursor ions was set to 10 ppm and the mass tolerance for fragment ions to 0.5 Da. Validation of MS/MS-based peptide and protein identification was performed with Scaffold V4.8.7 (Proteome Software, Portland, USA), and peptide identifications were accepted if they exhibited at least deltaCn scores of greater than 0.1 and XCorr scores of greater than 2.2, 3.3 and 3.75 for doubly, triply and all higher charged peptides, respectively. Protein identifications were accepted if at least 2 unique peptides were identified.

Volcano plot visualization of the mass spectrometry results was performed using Perseus (version 1.6.1.3)^69^ according to the following procedures. Contaminants and proteins with less than three valid values in at least one experimental group (Atpθ-3xFLAG and GFP-3xFLAG) were first removed from the matrix, and the normalized spectrum abundance factor (NSAF) intensities were log_2_-transformed. Imputation of the missing values was then performed based on the normal distribution of each column using default settings. Two sample *t* tests were performed before generating the final volcano plot. A heat map was generated by the hierarchical clustering function of Perseus 1.6.1.3 with default settings.

FLAG-tagged F_0_F_1_ ATP synthase was purified from *Synechocystis* 6803 overexpressing P*petE*::*atpB*-3xFLAG using a similar approach with modifications. *Synechocystis* 6803 overexpressing P*petE*::*atpB*-3xFLAG was cultured in Cu^2+^-free medium using the cell-DEG system in which much higher optical densities were obtained^52^. Cu^2+^ at 2 µM was added to the system when the OD_750_ reached 8.0 to induce the expression of the *petE* promoter, and the cells were collected by centrifugation (5,000 g, 10 min, room temperature). Cell pellets were washed once using prechilled FLAG buffer 2 (FLAG buffer without Tween-20). After disruption using Precellys (all remaining steps were performed at 4°C unless stated otherwise), the lysate was centrifuged at 4,000 g for 10 min to remove unbroken cells and beads, followed by 20,000 g for 1 h to collect the membrane fraction. The membrane fraction was then resuspended in FLAG buffer 2 supplemented with 1% *n*-dodecyl-beta-D-maltoside (β-DM) and incubated for 1 h with gentle agitation. Nonsoluble components were removed by centrifugation (20,000 g, 30 min), and the supernatant was filtered with a 0.45 µm syringe filter and then subjected to ANTI-FLAG M2 affinity agarose gel electrophoresis. The resin was prepared according to the manufacturer’s instructions using FLAG buffer 3 (FLAG buffer 2 supplemented with cocktail protease inhibitor and 0.03% [w/v] β-DM). The column was washed with 5 x 2 mL FLAG buffer 3 and eluted with 1.5 mL FLAG buffer 3 with 150 µg/mL 3xFLAG peptide (Sigma-Aldrich). The eluates were then concentrated to 200 µl with a 100 kDa MWCO centrifuge concentrator. The protein concentration was measured using the Bradford method.

#### RNA isolation and Northern blot

Cyanobacterial cells except those of *Nostoc* 7120 were harvested by vacuum filtration on hydrophilic polyethersulfone filters (Pall Supor®-800; 0.8 µm for *Synechocystis* 6803, *Thermosynechococcus elongatus* BP-1 and *Gloeobacter violaceus* PCC 7421, 0.45 µm for *Prochlorococcus* MED4). Total RNA was then isolated using PGTX^70^. The isolated RNA was mixed with 2x loading buffer (Ambion) and incubated for 5 min at 65°C. Denatured RNA samples were separated in a 1.5% agarose gel supplemented with 16% (v/v) formaldehyde and then transferred to a positively charged nylon membrane (Hybond™-N+, GE Healthcare) by capillary blotting with 20x SSC buffer (3 M NaCl, 0.3 M sodium acetate, pH 7.0) overnight.

After the RNA was cross-linked to the membrane by UV light (125 mJ), the membranes were hybridized with specific [γ-^32^P]ATP end-labeled oligonucleotides or [α-^32^P]UTP-labeled single-stranded RNA probes generated by *in-vitro* transcription from DNA templates using the MAXIscript® T7 In Vitro Transcription Kit (Ambion). The primers and oligonucleotides used for generating DNA templates are given in **Table S3**. Hybridization in 0.12 M sodium phosphate buffer (pH 7.0), 7% SDS, 50% deionized formamide and 0.25 M NaCl was performed overnight at 45°C or at 62°C with labeled oligonucleotide probes or labeled transcript probes, respectively. The hybridized membrane was then washed using washing solutions I (2xSSC, 1% SDS), II (1x SSC, 0.5% SDS) and III (0.1x SSC, 0.1% SDS) for 10 min each at 5 degrees below the hybridization temperature. Total RNA from *Nostoc* 7120 was prepared as described^71^ and separated on 8% urea-acrylamide gels. As a probe, a PCR fragment was generated as template to label one strand with Taq polymerase using only one oligonucleotide and [α-^32^]P-dCTP. Signals were visualized using Typhoon FLA 9500 (GE Healthcare) or Cyclone Storage Phosphor System (PerkinElmer) and Quantity One® software (Bio-Rad).

#### Membrane preparation and ATP hydrolysis assay

One liter of *Synechocystis* 6803 or *Thermosynechococcus elongatus* BP-1 cultures were grown to an OD_750_ of approximately 1 and cells were collected by centrifugation at 6,000 g for 5 min. The pellet was then washed once with precooled buffer A (1.0 M betaine, 0.4 M d-sorbitol, 20 mM HEPES-NaOH, 15 mM CaCl2, 15 mM MgCl2, 1 mM 6-amino-*n*-caproic acid, and protease inhibitor cocktail; pH 7.0) and then lysed using a Precellys homogenizer (steps afterwards were conducted at 4°C). Glass beads and unbroken cells were removed by centrifugation at 4,000 g for 10 min, and then the crude membranes were collected by centrifugation at 20,000 g for 1 h. The acquired membrane pellet was washed twice with buffer A, resuspended and incubated on ice for at least 1 h. Undissolved components were removed by centrifugation at 4,000 g for 5 min, and the membrane suspension was quantified by measuring the Chl *a* concentration at OD_664_^72^.

The ATPase activity of the membrane was measured via an ATP hydrolysis coupled enzyme activity assay. Buffer B (10 mM TES, 100 mM KCl, 1 mM MgCl2, and 0.1 mM CaCl2; pH 7.5) was supplemented with the indicated amounts of synthetic peptide or DCCD at room temperature. Then, 1 mM Mg-PEP, 0.175 mM NADH, 65 U pyruvate kinase (PK) and 82.5 U lactate dehydrogenase (LDH) was added. Before the activity measurement, 1 mM MgATP (pH 7.5) solution was added and incubated for 1 min to remove residual ADP. Then, membrane preparations containing approximately 10 μg Chl *a* were added to each assay, and the OD340 was measured immediately and after 10 min of incubation at room temperature using quartz cuvettes. The ATPase activity was calculated accordingly at nmol ATP mg Chl *a*^-1^ min^-1^. For the measurement of ATPase activity of the isolated ATP synthase, a similar method was applied, and 20 μg protein was used for each assay.

#### Far Western blotting

Far Western blotting was performed as described previously^38, 73^ with modifications. After electrophoresis, proteins were transferred onto a PVDF membrane. Synthetic AtpΘ peptide, anti-AtpΘ antiserum and anti-rabbit IgG antiserum were used for incubation sequentially. Milk powder was omitted in the denaturing/renaturing steps of the blotted membrane as described by Krauspe *et al*.^73^. The membrane with renatured proteins was first blocked with 5% milk powder in TBS-T and then incubated with 3 μg/mL synthetic AtpΘ peptide at 4°C overnight. Signals were detected with ECL start Western blotting detection reagent (GE Healthcare) on a chemiluminescence imager system (Fusion SL, Vilber Lourmat).

### QUANTIFICATION AND STATISTICAL ANALYSIS

Statistical analyses were performed with GraphPad Prism 6.0 (GraphPad Software, Inc., San Diego, CA). The ATP hydrolysis activities of membrane fractions isolated from different strains or conditions were compared using unpaired *t*-test with Welch’s correction (**Figure 5A** and **Figure S3; Data S1A** and **S1F**), and those of membranes isolated from the same strain but with different additives were compared using ratio paired *t*-test (**Figures 5A, 5B, 5C, 5D** and **Figure 6D; Data S1A, S1B, S1C, S1D** and **S1E**). Differences between groups were considered to be significant at a *P* value of <0.05 and very significant at a *P* value of <0.01.

### KEY RESOURCES TABLE

| REAGENT or RESOURCE | SOURCE | IDENTIFIER |
| --- | --- | --- |
| **Antibodies** |  |  |
| Mouse anti-FLAG M2 antibody | Sigma-Aldrich | Cat#A8592-1MG |
| Polyclonal AtpB antiserum \| Beta subunit of ATP synthase (chloroplastic + mitochondrial; rabbit antibodies) | Agrisera, Sweden | Cat# AS05 085-10 |
| Anti-Atpθ antiserum (rabbit antibodies) | Pineda Antibody-Service | this paper |
| Goat anti-rabbit secondary antibody | Sigma-Aldrich | Cat#A8275 |
| **Bacterial and virus strains** |  |  |
| *Synechocystis* sp. PCC 6803 | Cyanolab | Strain PCC-M |
| *Synechocystis* sp. PCC 6803 ($P_{atpT^-atpT}$) | This study | N/A |
| *Synechocystis* sp. PCC 6803 ($P_{atpT^-atpT^-3xFLAG}$) | This study | N/A |
| *Synechocystis* sp. PCC 6803 (Atpθ-KO) | This study | N/A |
| *Synechocystis* sp. PCC 6803 ($P_{petE^-sfGFP}$) | This study | N/A |
| *Thermosynechococcus elongatus* BP1 | Gen Enomoto | N/A |
| *Prochlorococcus* sp. MED4 | Claudia Steglich | N/A |
| *Gloeobacter violaceus* PCC 7421 | Mai Watanabe | N/A |
| *Nostoc* sp. PCC 7120 | Instituto de Bioquímica Vegetal y Fotosíntesis | N/A |
| *Nostoc* 7120 (GFP-less control) | This study | N/A |
| *Nostoc* 7120 ($P_{atpT^-gfp}$) | This study | N/A |
| *Nostoc* 7120 ($P_{atpT^-Atp\theta-GFP}$) | This study | N/A |
| **Chemicals, peptides, and recombinant proteins** |  |  |
| Atpθ peptide | JPT Peptide Technologies GmbH | this paper |
| AcnSP peptide | JPT Peptide Technologies GmbH | de Alvarenga et al. (2020)<sup>33</sup> |
| Atpθ_N peptide | JPT Peptide Technologies GmbH | this paper |
| Atpθ_C peptide | JPT Peptide Technologies GmbH | this paper |
| Atpθ_EE peptide | JPT Peptide Technologies GmbH | this paper |
| Atpθ_H peptide | JPT Peptide Technologies GmbH | this paper |
| **Deposited data** |  |  |
| Mass spectrometry comparing co-immunoprecipitated proteins of Atpθ-3xFLAG and sfGFP-Atpθ (*n*=1) | ProteomeXchange | PXD020126 |
| Mass spectrometry comparing co-immunoprecipitated proteins of Atpθ-3xFLAG and sfGFP-Atpθ (*n*=3; biological replicates) | ProteomeXchange | PXD024905 |
| **Oligonucleotides** |  |  |
| See Table S3 |  |  |
| **Recombinant DNA** |  |  |
| See Table S2 |  |  |
| **Software and algorithms** |  |  |
| Quantity One | Bio-Rad | Version 4.6.6 |
| Confocal Software (LCS) | Leica | Version 2.61 |
| GraphPad Prism | GraphPad Software, Inc. | Version 6.0 |

#### Overview on Supplementary Tables

**Table S1.** Number of putative *atpT* homologs, lengths, isoelectric points and amino acid sequences of all predicted AtpΘ homologs in the listed cyanobacteria. An alternative start codon for the AtpΘ homolog in *Pseudanabaena* sp. PCC 7367 is marked in red.

**Table S2.** Plasmids and vectors used in this work.

**Table S3.** Primers and oligonucleotides used in this work. Nucleotides in boldface letters highlight the sequence of the T7 promoter.

**Table S4.** Mass spectrometry results of pull-down experiment. Subunits of the ATP synthase are marked in bold. “High” is written for the log_2_ fold change if the respective protein was not detected in either one or both of the controls (P_*atpT*_-*atpT* and P_*petJ*_-3xFLAG-*sfgfp*).

**Table S5.** Raw mass spectrometry results of the second pull-down experiment. The table lists all proteins identified together with their quantitative values. Proteins, which have not been identified in a given samples, are marked with n.d. (not detected). The normalized spectrum abundance factor (NSAF) is used for quantification. The higher abundant a protein is in the sample, the higher is its quantitative value. Protein abundance is also visualized by a color gradient from green (low abundance) over yellow to red (highly abundant).

**Data S1.** Details for the statistical analysis. (**A**) Details for the statistical analysis in **Figure 5A**. (**B**) Details for the statistical analysis in **Figure 5B**. (**C**) Details for the statistical analysis in **Figure 5C**. (**D**) Details for the statistical analysis in **Figure 5D**. (**E**) Details for the statistical analysis in **Figure 6D**. (**F**) Details for the statistical analysis in **Figure S3**.

