## Supplementary material for "AtpΘ is an inhibitor of F_0_F_1_ ATP synthase to arrest ATP hydrolysis during low-energy conditions in cyanobacteria": Figures S1 to S6 and Tables S2 and S3

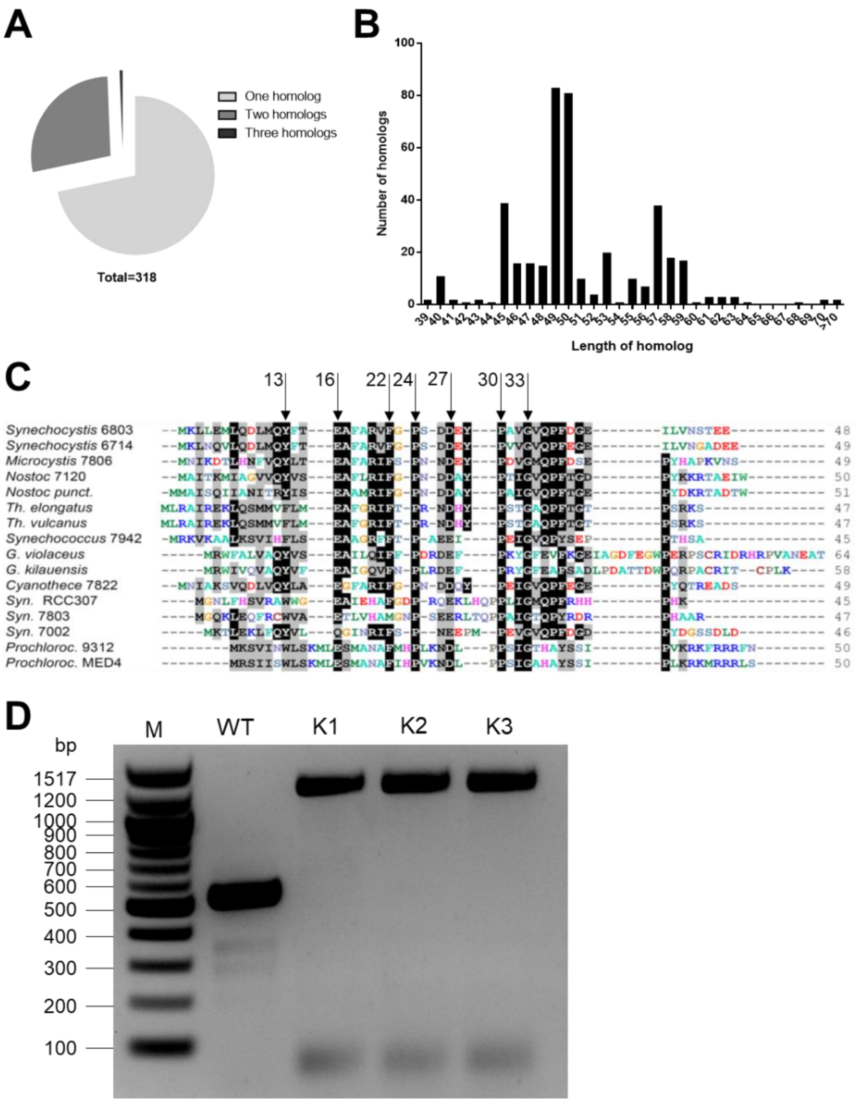


**Figure S1. Basic genetic and biochemical facts on *atpT* and AtpΘ. (A)** Pie chart of the number of cyanobacteria strains with one, two or three possible *atpT* homologs. **(B)** Length distribution of putative AtpΘ homologs. **(C)** Sequence comparison of AtpΘ homologs from *Synechocystis* 6803 and 6714, *Microcystis aeruginosa* sp. PCC 7806, *Nostoc* sp. PCC 7120 and *Nostoc punctiforme* sp. PCC 73102, thermophilic *Thermosynechococcus elongatus* BP-1 and *Thermosynechococcus vulcanus* NIES-2134*, Gloeobacter violaceus* sp. PCC 7421and *kilauensis* JS1*, Synechococcus* sp. PCC 7942, *Cyanothece* sp. PCC 7822, as well as the marine picocyanobacteria *Synechococcus* RCC307, WH 7803, WH 7002, *Prochlorococcus* MIT9312 and MED4. Widely conserved positions are shaded and the most conserved positions are labelled by vertical arrows and numbered according to the *Synechocystis* 6803 protein**. (D)** Genotype verification of *atpT* replacement by a chloramphenicol resistance cassette in *Synechocystis* 6803. Thirty-six cycles of colony PCR were performed with primers Norf1-KO-ver-F and Norf1-KO-ver-R (**Table S3**) flanking the gene. The calculated fragment length of 1244 bp was obtained for three independent clones of the mutant, while the amplicon obtained for the wild type matched the expected length of 541 bp. As size marker (M) the 100 bp DNA ladder (New England Biolabs, UK) was used.

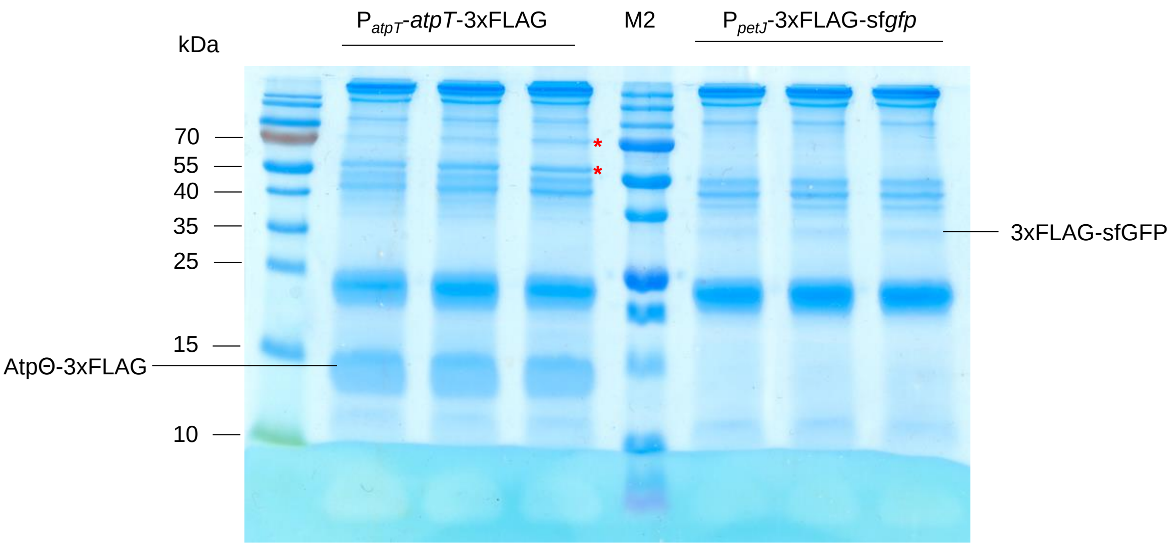


**Figure S2. Tricine SDS-PAGE characterization of protein pull-down samples.** Proteins eluted from Anti-FLAG M2 Magnetic Beads were separated on a Tricine-SDS gel, then stained with Coomassie Brilliant Blue. Three biological replicates were included for both strains overexpressing $P_{atpT}$-*atpT*-3xFLAG or $P_{petJ}$-3xFLAG-sf*gfp*. The suspected bands for AtpΘ-3xFLAG and 3xFLAG-GFP are indicated and two candidate proteins that were enriched only by pull-down with AtpΘ-3xFLAG are labelled by red asterisks. PageRuler™ Prestained Protein Ladder (10–170 kDa, Fermentas; left) and Protein™ DualXtra (2-250 kDa, Bio-Rad; lane M2) were used as molecular mass markers.

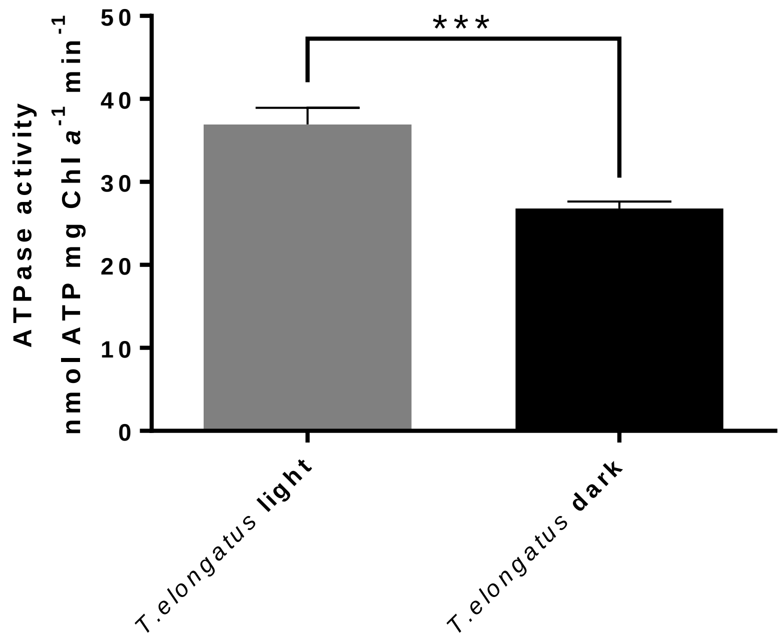


**Figure S3. Measurement of the ATPase activity in membrane fractions of *Thermosynechococcus elongatus* BP-1.** The membrane fraction was isolated from wild type *Thermosynechococcus elongatus* BP-1 cells grown under continuous light or after 24 hours of darkness incubation, then the ATPase activity was measured. Details for the statistical analysis can be found in **Data S1F**.

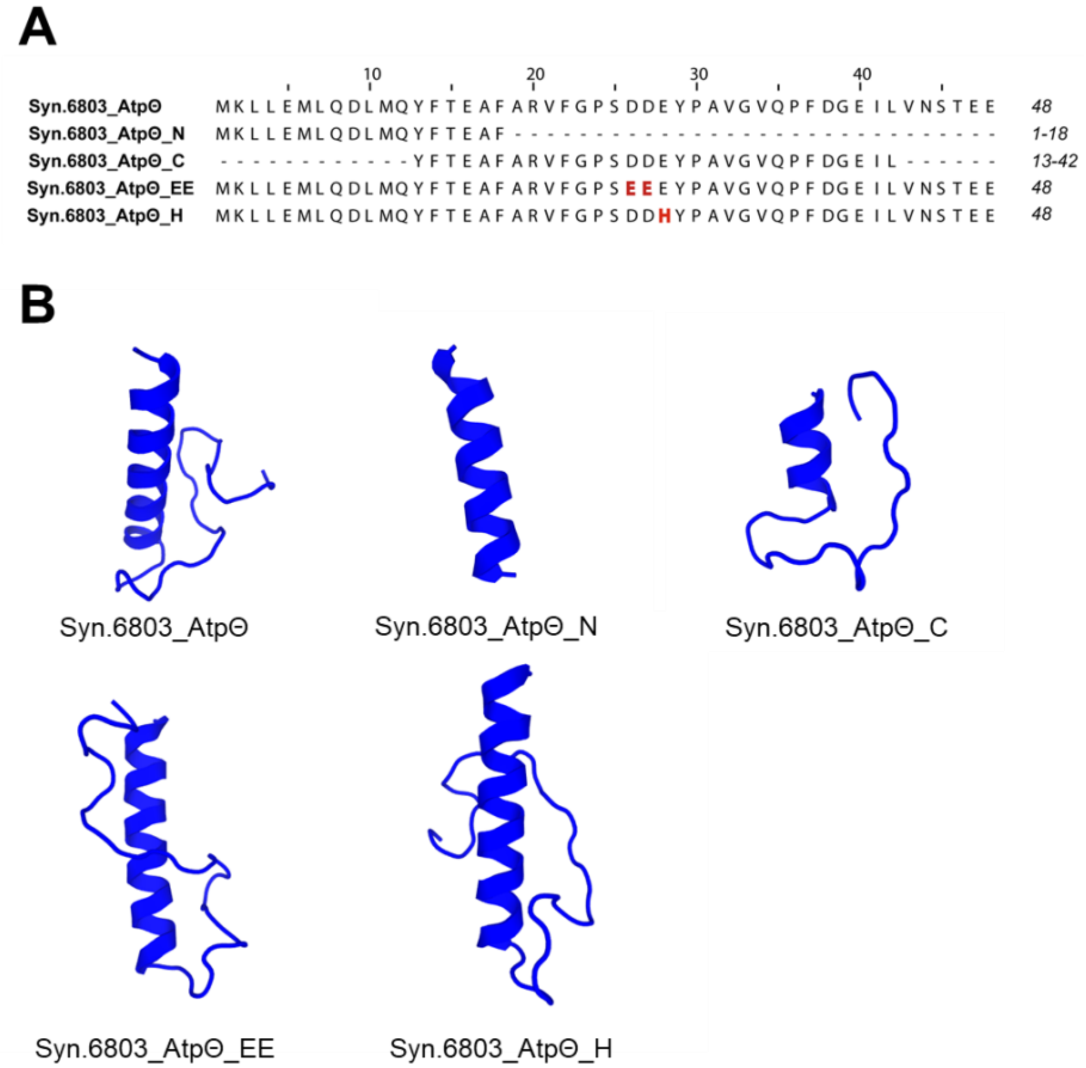


**Figure S4. Design and predicted structures of five *Synechocystis* 6803 AtpΘ mutant peptides. (A)** The amino acid sequences of the four mutant peptides. The substituted amino acids in AtpΘ_EE and AtpΘ_H are marked in red. **(B)** The predicted structure of *Synechocystis* 6803 AtpΘ and four mutant peptides using PEP-FOLD3[S1].

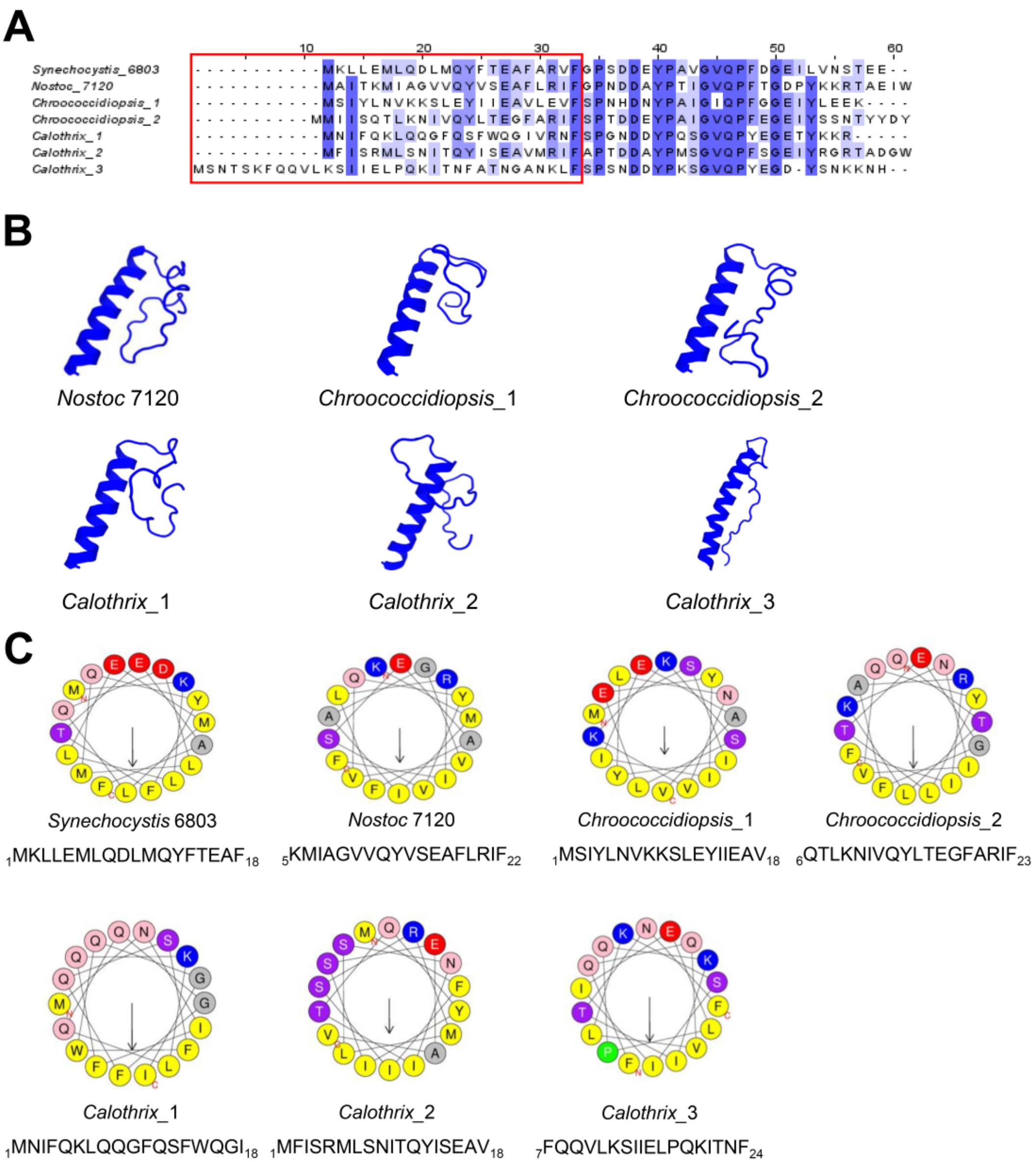


**Figure S5. Sequence alignment, predicted structures and amphipathic helices of AtpΘ homologs from cyanobacteria in which one, two and three homologs were identified. (A)** Alignment and predicted alpha-helical regions of AtpΘ from *Synechocystis* 6803 and six homologs from other cyanobacteria. The alignment was conducted using Jalview2 software. Conserved amino acids are marked with different shades of blue according to their percentage identity. The red box marks the position of the predicted N-terminal alpha helix. **(B)** Predicted structures of AtpΘ homologs in *Nostoc* 7120, *Chroococcidiopsis* sp. PCC 6712 (*Chroococcidiopsis*-1 and -2) and *Calothrix parasitica* NIES-267 (Calothrix-1, -2 and -3). All structures were predicted by PEP-FOLD3[S1] (four amino acids were omitted from the N terminus and C terminus of *Calothrix*-3 to keep the length limit of 50). **(C)** Predicted amphipathic helices within *Synechocystis* 6803 AtpΘ and the homologs in the same three other strains as in panel (B). The predicted unpolar stretches are indicated by arrows and the N and C termini are indicated. The prediction was performed using the HELIQUEST web server[S2].

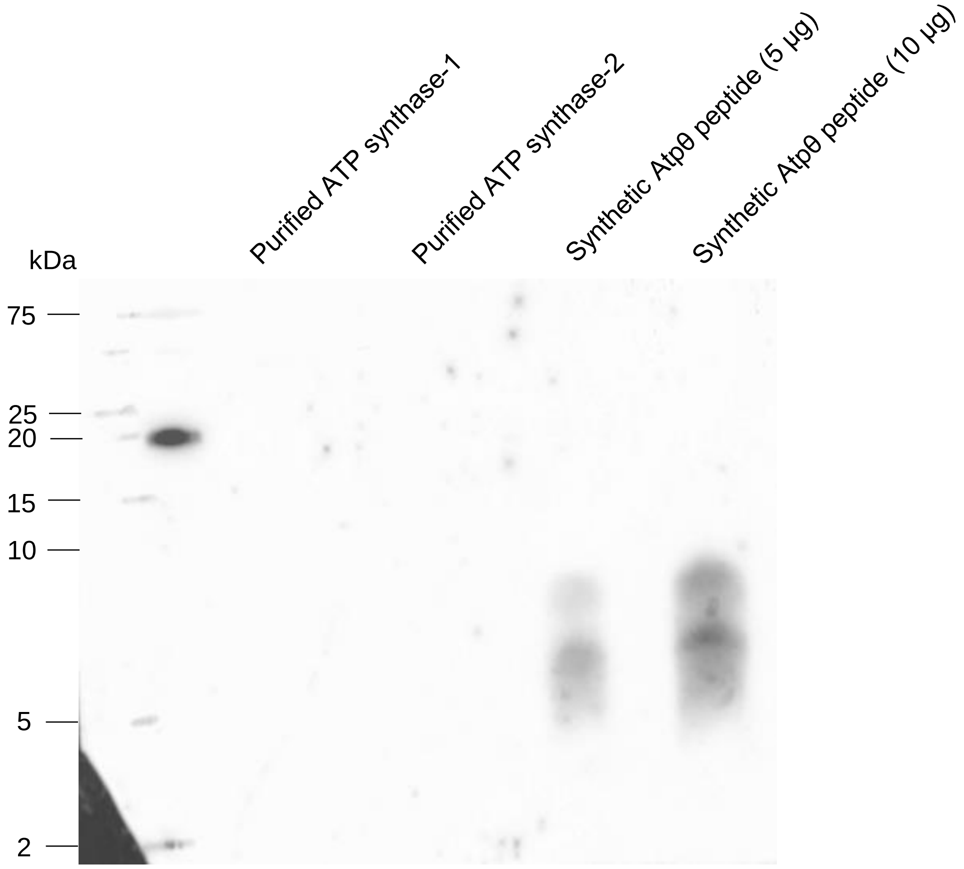


**Figure S6. Mock far Western blot characterization of the purified ATP synthase using anti-Atpθ antiserum.** 15 µg of two biological replicates of the purified ATP synthase samples were loaded on a tricine-SDS gel together with different amounts of synthetic Atpθ peptide as positive controls. The blotted membrane was incubated with TBST buffer instead of synthetic AtpΘ peptide before subjected to anti-AtpΘ antiserum and anti-rabbit antiserum. Precision Plus Protein DualXtra (2–250 kDa, Bio-Rad) served as molecular mass marker.

**Table S2**. Plasmids and vectors used in this work.

| ID | Comments | Reference |
|---|---|---|
| pCSEL24 | *gfp*-less control for *Nostoc* 7120 | Olmedo-Verd et al. (2006)[S3] |
| pSAM342 | Transcriptional fusion of DNA matching coordinates 2982431-2982070 from *Nostoc* 7120. PCR fragment amplified with primers 900+901, ClaI/XhoI-digested and cloned into *Cla*I/*Xho*I-digested pSAM270 [4]. | This work |
| pSAM344 | Translational Atpθ-GFP fusion for *Nostoc* 7120, coordinates 2982431-2981902. PCR fragment amplified with oligonucleotides 900+902, *Cla*I/*EcoR*V-digested and cloned into *Cla*I/*EcoR*V-digested pSAM147[S5] in frame with the gfpmut2 gene, rendering pSAM343. The *EcoR*I fragment from pSAM343, containing the fusion between the *atpT* promoter, the *atpT* gene and the gfpmut2 gene, was cloned into *EcoR*I-digested pCSEL24, yielding pSAM344. | This work |
| pUC-atpTKO-CmR | Replace the *atpT* gene in the genome of *Synechocystis* 6803 with a chloramphenicol resistance cassette (CmR) via homologous recombination. | This work |
| PatpT::atpT | Overexpressing AtpΘ with the native promoter of *atpT*. | Baumgartner et al. (2016)[S6] |
| PatpT::atpT-3xFLAG | Overexpressing FLAG-tagged AtpΘ with the native promoter of *atpT*. | |
| PpetJ::3xFLAG-sfgfp | Overexpressing FLAG-tagged sfGFP with the $P_{petJ}$ promoter. | |
| PpetE::atpB-3xFLAG | Overexpressing FLAG-tagged AtpB with the $P_{petE}$ promoter. | This work |

**Table S3.** Primers and oligonucleotides used in this work.

| Code | Sequence | Purpose |
|---|---|---|
| Norf1-KO-ver-F | GCCATAATCCACCATAGGTGAACC | Genotype verification of *atpT* deletion and replacement with a CmR cassette in *Synechocystis* 6803 (**Fig. S1**) |
| Norf1-KO-ver-R | GTGGTAAATGCCTGAGGAGATATGC | |
| #350 | TGATCGCTGGCGTAGTACAG | Template for *atpT* probe in *Anabaena* 7120 |
| #351 | ATTGATGAGTTCGTAGTAGCAGT | |
| #900 | GTTTTATCGATACAAGCAAATACAAGCTTTATAG | GFP fusions in *Anabaena* 7120 |
| #901 | ACTCCTACTCGAGTCAATTTCTG | |
| #902 | GTTTGATATCCCAAATCTCTGCTGTAC | |
| AtpTKO-up-F | CAAGGCGATTAAGTTGGGgctgaccatcttccctggattg | Amplification of *atpT* upstream flanking region |
| AtpTCmKO-up-R | GTGCCGATCAACGTCTCgtaaacagtactgaagaatagtcgatgg | |
| AtpTCmKO-down-F | GCCTGGTGCTACGCCTGgtaatttcatcataacctccgtcccaatg | Amplification of *atpT* downstream flanking region |
| AtpTKO-down-R | GAATTCGAGCTCGGTACgttcatcgagatgctcaccaatactgc | |
| AtpTKO-vec-F | GTACCGAGCTCGAATTCGTAATCATG | Amplification of pUC19 vector backbone |
| AtpTKO-vec-R | CCCAACTTAATCGCCTTGCAGCAC | |
| CmR-F | GAGACGTTGATCGGCACGTAAG | Amplification of chloramphenicol resistance cassette from pVZ321 vector |
| CmR-R | CAGGCGTAGCACCAGGCGTTTAAG | |
| Probe_atpT_fw | **GTAATACGACTCACTATAGGGAGA**CCATCGACTATTCTTCAGTACTGTTTAC | Amplification of template for *atpT* probe in *Synechocystis* 6803 |
| Probe_atpT_rev | TTGAGATGCTACAGGACCTTATGC | |

| | | |
|---|---|---|
| 5S Oligo PCC6803 | CTTGGCATCGGACTATTGTGCCGT | RNA quantification in *Synechocystis* 6803 |
| pUC19-XbaI_PpetE_fw | GCTCGGTACCCGGGGATCCTCTAGACTGGGCCTACTGGGCTATTC | Amplification of P*petE* introducing X*baI* site and creating overlaps with pUC19 and atpB |
| atpB::PpetE_rev | CGGCTACCATACTTCTTGGCGATTGTATCTATAGG | |
| PpetE::atpB_fw | GCCAAGAAGTATGGTAGCCGTAAAAGAAGCAAC | Amplification of *atpB* generating overlaps with P*petE* and 3xFLAG::3´UTR *atpT*::T*oop* |
| 3xFlag_atpB_rev | TATAATCCATACCCTCTTTGAGCTTGGCAC | |
| atpB_3xFlag_fw | CAAAGAGGGTATGGATTATAAAGATCATGATGGCGATTATAAAG | Amplification of 3xFLAG::3´UTR *atpT*::T*oop* introducing P*stI* site and creating overlaps with pUC19 and *atpB* |
| 3xFlag_PstI-pUC19_rev | GCCAAGCTTGCATGCCTGCAGAATAAAAACGCCCGGCGGC | |
| Probe_atpTGV_fw | **GTAATACGACTCACTATAGGGAGA**TCAGGTGGCTTCGTTTGCGACC | Template for *atpT* probe in *Gloeobacter violaceus* PCC 7421 |
| Probe_atpTGV_rev | ATGCGGTGGTTTGCTCTGGTGG | |
| Probe_GV_cont | CCTCCAAGTATCTTCGCTGCTGC | RNA quantification in *Gloeobacter violaceus* PCC 7421 |
| Probe_BP1_fw | **GTAATACGACTCACTATAGGGAGA**CTTGCGGAAGAGGGTCACTAGCTC | Template for *atpT* probe in *Thermosynechococcus elongatus* BP-1 |
| Probe_BP1_rev | GAGCAATACGTGAGAAACTGCAAAGTATG | |
| Probe_BP1_cont | GAGTTCGGGATGGATCAGCGTG | RNA quantification in *Thermosynecho-* |

| | | |
|---|---|---|
| | | *coccus elongatus* BP-1 |
| Probe_MED4_fw | **GTAATACGACTCACTATAGGGAGA**CTAGCTGAGTCTTCTTCGCATTTTCC | Template for *atpT* probe in *Prochlorococcus marinus* MED4 |
| Probe_ MED4_rev | CAAATGAGATCAATTATTAGTTGGCTTTC | |
| Probe_ MED4_cont | GTTCGAGATGGATCGGAGTGGTTC | RNA quantification in *Prochlorococcus marinus* MED4 |
